# Cortico-thalamic communication for action coordination in a skilled motor sequence

**DOI:** 10.1101/2023.10.25.563871

**Authors:** Yi Li, Xu An, Patrick J. Mulcahey, Yongjun Qian, X. Hermione Xu, Shengli Zhao, Hemanth Mohan, Shreyas M. Suryanarayana, Ludovica Bachschmid-Romano, Nicolas Brunel, Ian Q. Whishaw, Z. Josh Huang

**Affiliations:** Department of Neurobiology, Duke University, Durham, NC 27710, USA; Cold Spring Harbor Laboratory, Cold Spring Harbor, NY 11724, USA; Department of Biomedical Engineering, Duke University, Durham, NC 27710, USA; Department of Neuroscience, Canadian Centre for Behavioural Research, University of Lethbridge, Lethbridge, AB, T1K 3M4, Canada

## Abstract

The coordination of forelimb and orofacial movements to compose an ethological reach-to-consume behavior likely involves neural communication across brain regions. Leveraging wide-field imaging and photo-inhibition to survey across the cortex, we identified a cortical network and a high-order motor area (MOs-c), which coordinate action progression in a mouse reach-and-withdraw-to-drink (RWD) behavior. Electrophysiology and photo-inhibition across multiple projection neuron types within the MOs-c revealed differential contributions of pyramidal tract and corticothalamic (CT^MOs^) output channels to action progression and hand-mouth coordination. Notably, CT^MOs^ display sustained firing throughout RWD sequence and selectively enhance RWD-relevant activity in postsynaptic thalamus neurons, which also contribute to action coordination. CT^MOs^ receive converging monosynaptic inputs from forelimb and orofacial sensorimotor areas and are reciprocally connected to thalamic neurons, which project back to the cortical network. Therefore, motor cortex corticothalamic channel may selectively amplify the thalamic integration of cortical and subcortical sensorimotor streams to coordinate a skilled motor sequence.

## INTRODUCTION

Animals deploy skilled motor behaviors involving the orderly coordination of multiple movements across the body to achieve ethological goals. For example, consummatory behaviors in rodents and primates often consist of reaching for a food item with the forelimb, grasping food with the hand, and withdrawing the hand to the mouth to eat or drink^1,2^. These elemental actions are sequentially executed and continuously coordinated with concurrent sensory streams to compose a skillful and goal-directed consumption behavior. The view of behavior as involving dynamic unfolding of serially ordered and coordinated constituent movements was championed by Lashley^3^ decades ago, but the underlying brain circuit mechanisms remain poorly understood.

The individual actions of reach, grasp, and lick can be elicited from specialized spinal and brainstem motor centers^4–6^. The orderly progression and coordination of these actions to compose a complex behavior likely involve inter-regional communications that integrate motor-related signals with sensory feedback to refine movement commands during the action sequence^7–10^. Across the different levels of motor control infrastructure, the cerebral cortex comprises a constellation of functional areas that integrate motor plan with cognitive and multi-sensory information, and broadcast the outcome of cortical processing to multiple subcortical sensorimotor centers^10–15^. Decades of studies have examined various motor areas in controlling individual actions such as the reach^2,16,17^ and lick^18–21^. However, how cortical circuits and their constituent cell types and output channels regulate the coordination of forelimb and orofacial movements to compose a complex ethological behavior remains an open question.

Among the major cortical glutamatergic projection neuron (PN) classes, whereas the intratelencephalic (IT) PNs assemble intra-cortical and cortico-striatal processing streams, the extratelencephalic (ET) PNs constitute multiple parallel output channels that communicate with myriad subcortical regions^12,22,23^. Of the two major ET neuron types, the pyramidal tract (PT) PNs project to multiple structures, from the basal ganglia to the spinal cord, thus broadcasting cortical signals brain-wide^10,12,23–25^. The corticothalamic (CT) PNs project exclusively to the thalamus^22,23,26^, a central hub that integrates multi-sensory, motor, and body state information and in turn influences ongoing cortical network activity^27–30^. Whether and how these distinct output channels within specific cortical areas regulate action progression and coordination during a complex motor sequence is unclear; in particular, the role of the corticothalamic pathway is largely unexplored.

Here, combined quantitative behavior analysis, wide-field imaging and optogenetic manipulation uncovered a cortical network and one of its key nodes, the central region of the secondary motor cortex (MOs-c) that facilitates the orderly progression and coordination of a reach-and-withdraw-to-drink (RWD) behavior. Subsequent PN-type resolution electrophysiology and optogenetic inhibition in the MOs-c revealed the activity dynamics of two distinct cortical output channels, the *Fezf2*-expressing PT (PT^Fezf2^) and *Tle4-* expressing CT (CT^Tle4^) neurons, and their differential contribution to RWD progression and coordination. Notably, CT^Tle4^ manifested sustained dynamics across RWD action phases and amplified similarly sustained activities in their postsynaptic thalamus neurons, which also contributed to action progression and coordination. MOs-c CT^Tle4^ received converging inputs from forelimb and orofacial sensorimotor areas of the RWD network and were reciprocally connected to thalamic neurons, which in turn projected back to this cortical network. Our findings highlight the key and unexpected role of corticothalamic communication in a high-order motor cortex in facilitating action progression and coordination in a skilled sequential motor behavior.

## RESULTS

### Reach-to-consume involves the ordering and coordination of multiple forelimb and oral actions

The reach and withdraw to drink (RWD) task was performed in head-restrained mice (**Fig. 1a**). In this task, mice use chemosensory and vibrissae cues to locate a waterspout in the dark with which to guide their left forelimb to grasp a water drop that they then withdraw to the mouth to drink by licking the hand^31^. Combining high-speed videography and deep neural network-based behavior tracking^32^ (**Fig. S1a-i**), we extracted thirteen movement time series of the left hand and its relationship with the waterspout, mouth, and other body parts (**Fig. S1j**, see Methods). Dimensional reduction analysis of these feature time series revealed three major action phases, reach, withdraw, and consume that constitute the full behavior (**Fig. S1k**). The *reach* involves a mouse *aiming* its hand after lifting it with the digits closed and flexed, *advancing* the hand with combined hand supination and digits extension towards the target to *grasp* the water (**Fig. 1b-c, Supplementary video 1**). The *withdraw* involves the hand supinating to bring water to the mouth with a further supination and finger opening to release the water. The mouth then opens, and the tongue protrudes to *lick and consume* the water from the hand; the hand remains supinated close to the mouth during the multiple subsequent licks that consume the water (**Fig. 1b-c, Supplementary video 1**). Mice usually lift the hand shortly after water delivery (429.2 ± 478.2 ms, median ± SD, 6392 trials) (**Fig. S1l**); the time course of the behavior is fast and varies trial by trial (516.7 ± 550.0 ms) comprising 308.3 ± 388.6 ms reach and 175.0 ± 354.4 ms withdraw (**Fig. 1d, S1m**). Tongue protrusion probability before water grasping is low (10% ± 13%) (**Fig. S1n**). The withdraw brings the hand close to the mouth with a further supination that is coordinated with tongue protrusion for licking (**Fig. 1e**). These results suggest that RWD involves the orderly spatial and temporal coordination of multiple movements across the reach, grasp, withdraw and hand lick, including digit opening during reach in anticipation of grasp, and hand supination toward the mouth during withdraw that is coordinated with tongue protrusion.

**Fig 1.**
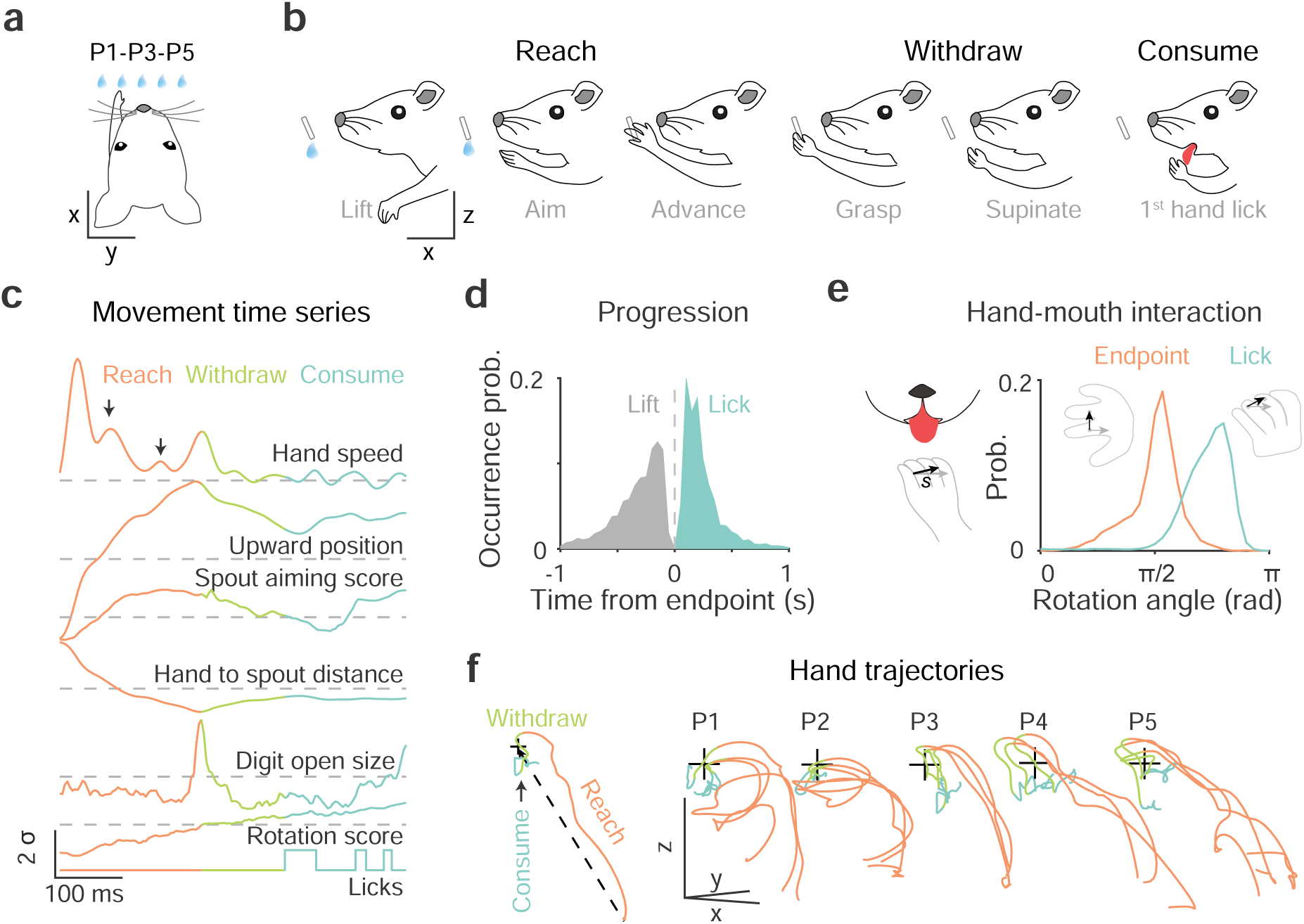
Coordinated progression of reach and withdraw to consume (RWD) sequence. **a.** Schematic of a head-restrained mouse reaching with left hand for a waterspout positioned at one of five locations (ipsilateral P1, P2; central P3; contralateral P4, P5). **b.** RWD involves reach, withdraw and consume action phases. The reach begins with the hand lift from the start position to the advance endpoint at the waterspout; withdraw starts with the grasp at the reach endpoint and ends with licking onset; consume includes licking and ends when the hand is replaced at the start position. **c.** Movement time series of reach onset to consume. The horizontal scale is time, and the vertical scale shows movement kinematics (σ). Arrows point to two separate hand speed changes that reflect sub-movement adjustments during reaching. The color code for reach, withdraw and consume is used in all subsequent figures. **d.** Distributions of the occurrence of onset timepoints of the reach and the first hand lick relative to the reach endpoint (0, dashed vertical line). (Reach duration: 308.3 ± 388.6 (median ± SD) ms; withdraw duration: 175.0 ± 354.4 ms; *n* = 6392 trials of 74 sessions from 27 mice.) **e.** Hand-mouth coordination upon the onset of tongue protrusion to lick water from the hand. The rotation angle is the direction change of the hand rotation vector (***s***) from the resting posture, at which ***s*** is in the opposite direction of the horizontal reference vector (gray). Rotation angle 0 and π indicate palm facing downward and upward, respectively. Endpoint: 1.67 ± 0.35 rad; lick onset: 2.38 ± 0.38 rad. *n* = 6392 trials. **f.** Waterspout dependent modulation of hand trajectories. Three random trials were annotated with action phases for each waterspout position in the same session. The dashed line in the schematic inset indicates the reference direction in relation to waterspout location (+).

To examine the effect of variation of target location on RWD movement, we presented the waterspout at five locations at random (**Fig. 1a, 1f**): a central location (P3) aligned to the nose, two ipsilateral locations on the same side of the reaching left hand (P1, P2), and two contralateral locations (P4, P5). After training, mice retrieved water with accurate reach endpoints at which the hand fully opened to grasp the water regardless of the target’s changing locations (**Fig. S1o**). In doing so, mice engaged different forelimb trajectories that resulted from changing spatiotemporal coordination of the arm, wrist, and digit movements during the process (**Fig. 1f, S1p**). With changing target location, the mouse adjusts its aim by upper arm abduction or adduction and wrist flexion to point the digits to the target location. This aiming phase during contralateral reaches occurs farther from the waterspout and requires larger angular corrections than ipsilateral reaches (**Fig. S1q-r**). The advance further adopts the palm-facing direction with digits opening and extension relative to target, and both the aim and advance often involve adjustments reflected in the speed of the reaching hand (**Fig. 1c, S1s**). Together, these results indicate that reaching is not a ballistic movement but rather involves the orderly coordination of arm, hand, and digit movements in relation to sensory features of the target, and withdraw-to-consume involves the coordination of forelimb and oral actions.

### Cortical subnetwork dynamics reports RWD progression

Leveraging multiple mouse driver lines targeting GCaMP6 expression to a set of genetic and projection defined PN subpopulations, we surveyed neural dynamics across the dorsal cortex of mice performing RWD with wide-field calcium imaging^33,34^ (**Fig. 2a,b, S2a**). Among these, the *Emx1-Cre* line targets most if not all PNs, *Cux1-CreER* targets L2/3/4 IT neurons that project largely within the cortex, *PlxnD1-CreER* targets L2/3/5a IT subset with strong projection to cortex and striatum, *Fezf2-CreER* targets L5 PT that project to the spinal cord, brainstem, thalamus, and basal ganglia structures, and *Tle4-CreER* targets L6 CT neurons that project almost exclusively to the thalamus^22^ (**Fig. S2b-c**). These PN subpopulation driver lines facilitate the detection of neural activity that might be masked in whole-population recording.

**Fig 2.**
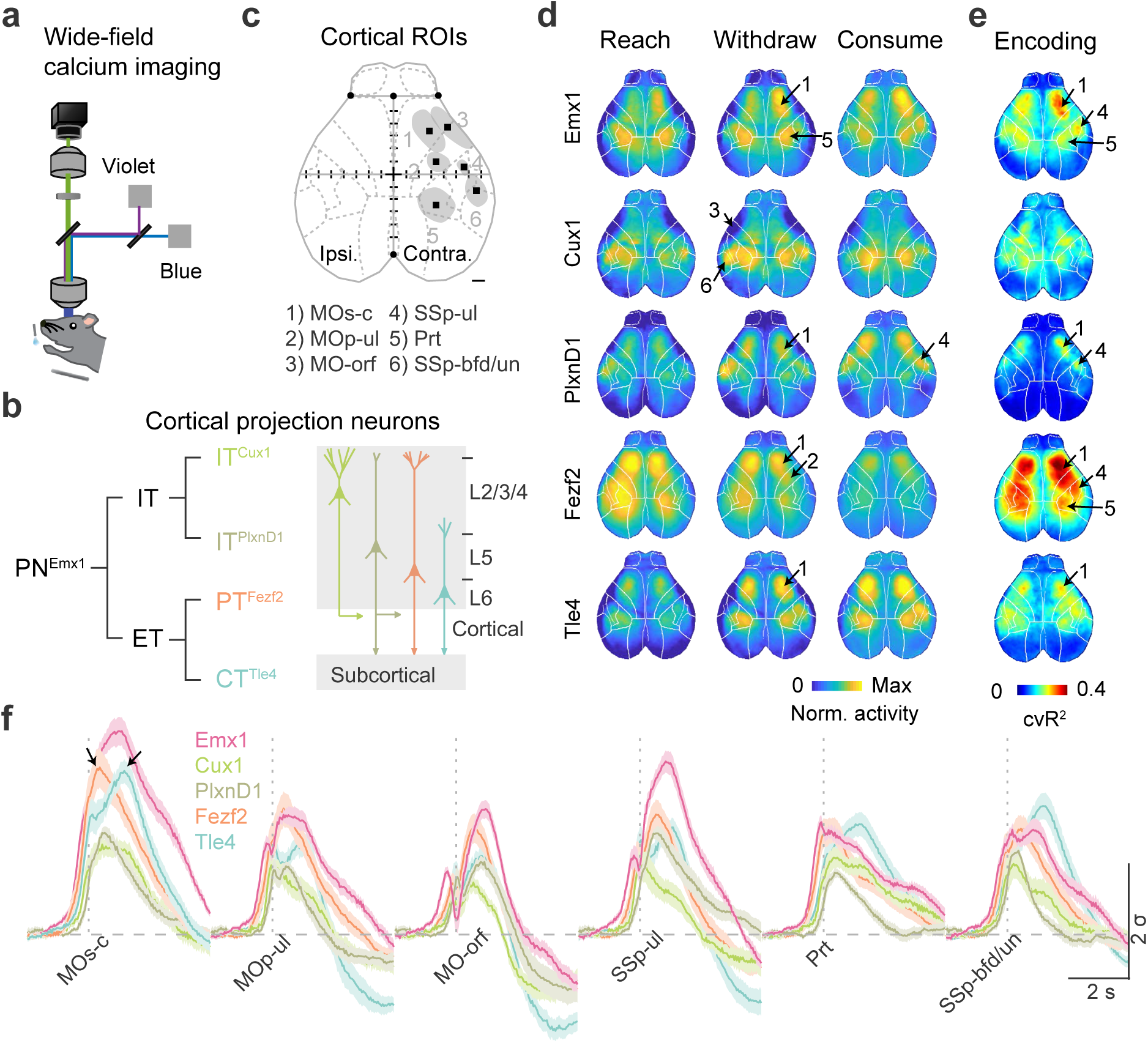
Cortical subnetworks differentially report movement progression. **a.** Schematic of wide-field calcium imaging for head-restrained mice reaching with the left forelimb. A violet channel was used as the control for the GCaMP channel (blue) to extract calcium activity. sCMOS camera, scientific complementary metal–oxide–semiconductor; LED, light-emitting diode. **b.** Genetically- and projection-defined projection neuron (PN) types (left) and their layer (L) distribution (right). The intratelencephalic (IT) class includes IT^Cux1^ and IT^PlxnD1^ types; the extratelencephalic (ET) class includes the pyramidal tract (PT^Fezf2^) and corticothalamic (CT^Tle4^) types. Emx1 marks all PNs. **c.** The cortical regions of interest (ROIs) contralateral to the reaching hand isolated from calcium fluorescence change during RWD. 1) MOs-c: secondary motor cortex central region; 2) MOp-ul: forelimb primary motor cortex; 3) MO-orf: orofacial motor cortex; 4) SSp-ul: anterior-lateral forelimb somatosensory cortex; 5) Prt: parietal cortex; 6) SSp-bfd/un: anterior part of barrel field and the unassigned region. Dashed lines indicate the boundaries of brain regions registered to Allen CCF. Homotypic ROIs on the ipsilateral hemisphere are not shown. Black squares indicate the center of each region. +, Bregma. Scale, 0.5 mm. **d.** PN-type specific cortex-wide calcium activity changes during reach, withdraw and consume. Data was averaged across all trials. Arrows point to the 6 ROIs. Cortex-wide calcium activity was registered to the Allen CCF and normalized to the max activity change. (*n* = 9 sessions from 5 mice for PN^Emx1^; 7 sessions from 4 mice for IT^Cux1^; 11 sessions from 4 mice for IT^PlxnD1^; 12 sessions from 6 mice for PT^Fezf2^; 10 sessions from 5 mice for CT^Tle4^. The same animals were used in subsequent panels.) **e.** Performance of generalized linear encoding models (cross-validated variance explained, cv*R*^2^), in which the movement time series of the reaching forelimb was used to predict calcium activity change. Warmer color indicates higher performance in explaining activity with forelimb movement. **f.** Calcium activity change of contralateral ROIs aligned to waterspout contact (vertical gray dash line) for different PNs during RWD from P2. σ: standard deviation. Arrows indicate “fast” and “delayed” activity peaks of PT^Fezf2^ and CT^Tle4^, respectively. Error shading: SEM.

Following training at a fixed P2 location with the left hand (**Fig. S2d**), we observed widespread activity dynamics bilaterally in all five PN populations during RWD (**Fig. S2e**). Cortex-wide activity patterns in all five PN populations changed with successful RWD progression (**Fig. 2d**) and were tightly correlated with target location (**Fig. S2f**). To extract the network dynamics correlated with moment-to-moment motor progression, we built a generalized linear encoding model (GLM) considering ten forelimb movement time series to predict relative calcium fluorescent fluctuations (**Methods**). The model performance, as reflected by 10-fold cross-validated variance explained (cv*R*^2^ value), reveals correlation strength between movement time series and normalized activity dynamics. A summary of GLM performance from all PNs revealed three activity nodes that were correlated with forelimb movements (**Fig. 2c, S2g;** see **Methods)**: the central region of the secondary motor cortex (MOs-c, partially overlapping with the rostral forelimb area (RFA)^35^, the forelimb somatosensory area (SSp-ul), and the parietal area (Prt). Analysis of IT^Cux1^ and IT^PlxnD1^ calcium signals additionally revealed nodes related to the anterior whisker barrel/unassigned sensory (SSp-bfd/un) and orofacial motor cortex (MO-orf)^36^. Forelimb primary motor area (MOp-ul)^23^ was isolated by comparing left and right hemisphere PT^Fezf2^ activity. With some regional differences, we observed higher correlations with RWD progression for PN^Emx1^, PT^Fezf2^, and CT^Tle4^ populations than for IT^Cux1^ and IT^PlxnD1^ populations (**Fig 2e, S2i**). Notably, the MOs-c CT^Tle4^ show a strong but “delayed” activation peak compared with the “fast” peak of PT^Fezf2^ (**Fig. 2f, S2h**). These results identify a cortical subnetwork with PN type resolution (**Fig. 2c**) in which PT^Fezf2^ and CT^Tle4^ activities are differentially correlated with the forelimb action phase progression.

### MOs-c is required for RWD action progression

To examine the behavioral role of each of the network nodes (**Fig. 2c**), we used closed-loop photoinhibition by activating parvalbumin (Pvalb) inhibitory interneurons in each of the identified cortical areas in *Pvalb;Ai32* mice (**Fig. S3a**). Inhibition was triggered during reach with a latency of 129.2 ± 42.8 ms (mean ± SD) upon hand lift. Among the tested areas, inhibition of MOs-c contralateral to the reaching hand resulted in deficits in a number of movement features, including a decreased probability of reach, withdraw, and consume (**Fig. 3a, S3b-c**). Inhibition of SSp-ul resulted in deficits in digit closing, supination, and hand lick (**Fig. S3b-c**), suggesting that this forelimb somatosensory area may contribute to action coordination and refinement. Cortical inhibition ipsilateral to the reaching hand did not produce significant effects (**Fig. S3b-c**). We then analyzed how MOs-c suppression interfered with RWD action progression in relation to reach, withdraw and consume. In about 30% of the trials, inhibition following hand lift resulted in a collapse of the reaching movement (**Fig. 3b**). In trials when the hand did reach the target, supination during withdraw and/or hand lick often failed to occur (**Fig. 3c, S3d-e**). Upon termination of inhibition, the mouse immediately resumed and completed the RWD act (**Fig. S3f**), consistent with previous observations^2,31^. In summary, amongst the contralateral cortical network nodes, the inhibition effect of MOs-c makes it stand out for its crucial contribution to the RWD sequence.

**Fig 3.**
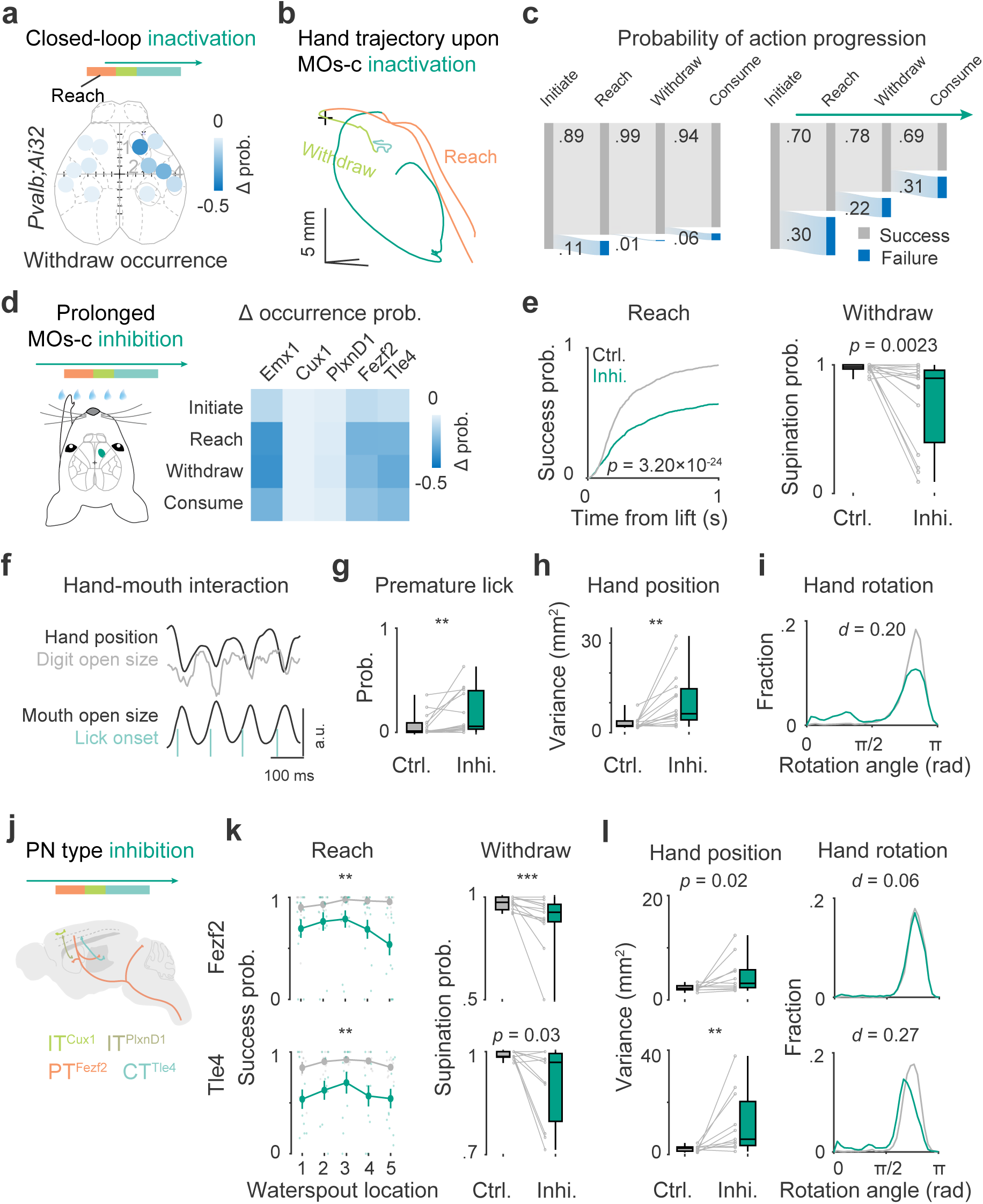
MOs-c PT and CT are required for the progression and coordination of RWD. **a.** Photoinhibition mapping of cortical areas by closed-loop activation of inhibitory interneurons. Inactivation of the contralateral areas of MOs-c (1), MOp-ul (2), and SSp-ul (4) decreased supination probability. Color scale represents changes in success probability between inhibition and control trials. (*n* = 5 *Pvalb;Ai32* mice, see **Supplementary Table 1** for statistics) **b.** Perturbation of movement progression upon close-loop MOs-c inactivation during reach in an example control and inhibition movement trajectory. In the inhibition trial, the reach was aborted; the hand returned to the start position, followed by another failed attempt. **c.** Impaired action sequence progression in control (left) and inhibition (right) trials upon contralateral MOs-c inhibition. 18% (81/441) control trials and 63% (209/331) inhibition trials failed to complete the RWD sequence. (*n* = 5 *Pvalb;Ai32* mice) **d.** Effects of prolonged inhibition of PN types by expressing optogenetic inhibitory opsins in MOs-c (turquoise). Heatmap summarized the change in success probability of action phases upon prolonged inhibition of each PN type compared with control trials. Same mice for all following panels. (*n* = 15 sessions from 8 PN^Emx1^ mice; 8 sessions from 6 IT^Cux1^ mice; 12 sessions from 7 IT^PlxnD1^ mice; 13 sessions from 7 PT^Fezf2^ mice; 11 sessions from 6 CT^Tle4^ mice.) **e.** Reduction of reach and withdraw success probability with prolonged PN^Emx1^ inhibition. Reach was quantified as target contact probability from all trials with successful lifts (two-sample Kolmogorov-Smirnov (KS) test, KS distance (*d*) = 0.72, *p* = 3.20×10^-24^). Withdraw was quantified as supination probability with successful reach (Wilcoxon rank sum test, \*\**p* < 0.01). **f.** Coherent hand-mouth movement time series during drinking. Hand upward position, digit open size, mouth open area, and lick onset variables are indicated. **g.** Increased premature lick probability during PN^Emx1^ inhibition (*n* = 15, Wilcoxon rank sum test, \*\**p* < 0.01). **h.** Increased variance in hand position relative to the mouth upon the onset of hand lick during PN^Emx1^ inhibition (*n* = 15, Wilcoxon rank sum test, *p* < 0.001). **i.** Abnormal hand posture during drinking with prolonged PN^Emx1^ inhibition indicated by palm-facing direction at lick onset (*n* = 13734 control and 9186 inhibition licks, two-sample KS test, *d* = 0.20, *p* = 1.61×10^-202^). **j.** Schematic of the prolonged inhibition of MOs-c PN types. **k.** PT ^Fezf2^ and CT^Tle4^ inhibition on success probability of reach and withdraw. Left: ANOVA; PT^Fezf2^ inhibition *F*_1,56_ = 11.52, \*\**p* = 0.0053; CT^Tle4^ inhibition *F*_1,40_ = 10.82, \*\**p* = 0.0082. Note the target location-dependent impairment in PT^Fezf2^ (inhibition × target *F*_4,56_ = 4.41, *p* = 0.0041) but not in CT^Tle4^ (inhibition × target *F*_4,40_ = 0.51, *p* = 0.7280). Right: Wilcoxon rank sum test, \*\*\**p* < 0.001. **l.** Variance in hand position (left) and hand posture (right) during PT ^Fezf2^ and CT^Tle4^ inhibition. (Left: Wilcoxon rank sum test, \*\**p* < 0.01. Right: two-sample KS test, \*\*\**p* < 0.001.)

### MOs-c PT and CT neurons mediate RWD action progression and coordination

To dissect the neural communication streams within MOs-c that contribute to RWD, we systematically examined several major PN projection types. First, we examined the effects of inhibiting all PNs by expressing a light sensitive inhibitory opsin GtACR1^37^ in *Emx1-Cre* mice using a Cre-dependent AAV vector (**Fig. S4a**). Closed-loop inhibition of MOs-c PNs^Emx1^ resulted in similar impairments of motor progression as those obtained with the activation of MOs-c Pvalb interneurons (**Fig. S4a**). Prolonged inhibition of MOs-c PNs^Emx1^ spanning the entire trial only slightly decreased lift probability (**Fig. S4b**), suggesting a minor role of MOs-c in the initiation of RWD movement. On the other hand, consistent with closed-loop inhibition, prolonged inhibition of PNs^Emx1^ in contralateral MOs-c decreased the probability of movement progression at multiple stages of the reach and the withdraw to consume movements (**Fig. 3d**). In trials when the hand did reach the waterspout, we observed a decrease of withdraw (**Fig. 3e**) and multiple deficits in hand-mouth coordination during consumption (**Fig. 3f-i**). Indeed, prolonged inhibition resulted in uncoordinated hand-mouth movements during drinking, reflected as increased premature tongue protrusion before grasp (**Fig. 3g**), increased variation of hand position upon lick (**Fig. 3h**), abnormal hand posture (**Fig. 3i**), and decreased coherence between hand and mouth movements (**Fig S4c**). Altogether, these results suggest that MOs-c is not crucial for the execution of individual actions (e.g. lift and lick) but it is involved in the orderly progression and coordination of these actions, especially the coordination between hand and oral actions.

Next, we optogenetically inhibited multiple PN subpopulations by virally expressing inhibitory opsin GtACR1 in four driver lines, *Cux1-*, *PlxnD1-*, *Fezf2-,* or *Tle4-CreER*, respectively (**Fig. 3j, S4d**). We analyzed the prolonged inhibition effect on the progression and coordination of RWD constituent actions (**Fig. 3d**). IT^Cux1^ inhibition had no significant effect on RWD progression (mixed-design ANOVA, *p* = 0.81 for reach; **Fig. S4e**). IT^PlxnD1^ inhibition led to a mild increase of premature lick (Wilcoxon rank sum test, *p* = 0.03) but rarely disrupted RWD progression (mixed-design ANOVA, *p* = 0.30 for reach; **Fig. S4e**). On the other hand, PT^Fezf2^ and CT^Tle4^ inhibition resulted in multiple deficits in action progression and coordination, including decreased reach (**Fig. 3k**), withdraw (**Fig. 3k**), and decreased hand-mouth coordination for consumption (**Fig. 3l**). Whereas the deficits of PT^Fezf2^ inhibition were waterspout location dependent, i.e. more pronounced when reaching for more difficult contralateral locations (mixed-design ANOVA, inhibition × target *F*_4,56_ = 4.41, *p* < 0.01), CT^Tle4^ inhibition effects were less modulated by waterspout location (mixed-design ANOVA, inhibition × target *F*_4,40_ = 0.51, *p* = 0.73; **Fig. 3k**). Moreover, CT^Tle4^ inhibition led to significant deficit in hand-mouth coordination, as indicated by increased variance in hand position (Wilcoxon rank sum test, *p* < 0.01) and hand rotation at the time of tongue protrusion for drinking (**Fig. 3l**), which is stronger than that of PT^Fezf2^ inhibition. Together, these results reveal the differential contributions of PN subpopulations and the requirement of CT^Tle4^ and PT^Fezf2^ activity in the orderly progression of RWD action sequence. In particular, they highlight the role of cortico-thalamic communication mediated by CT^Tle4^ in the coordination of hand and mouth movements.

### MOs-c CT and PT dynamics differentially correlate with RWD progression

To explore the neural coding properties of individual MOs-c neurons during RWD sequence, we performed electrophysiological recordings with linear probes. Individual neurons exhibited diverse spiking patterns tightly coupled to RWD actions, with varying peak amplitude and latency relative to reach onset (**Fig. S5a-c**). Deep layer neurons tended to be more strongly correlated with forelimb movements (**Fig. S5d-e**). Simultaneously recorded MOs-c population activity significantly decoded a range of arm, hand, and orofacial movement time series (**Fig. S5f-g**). In addition, MOs-c activity decoded target locations even before hand lift and its accuracy in doing so became more accurate during subsequent actions (**Fig. S5h**). These results suggest that MOs-c population activity is closely correlated with the progression of RWD movement sequence.

By applying optogenetic tagging, we next examined how the spiking dynamics of different MOs-c PN subpopulations correlated with the unfolding of the RWD movement sequence. We delivered brief light pulses to MOs-c to evoke the spiking of ChR2-expressing neurons during recording, and further applied statistical tagging analysis with criteria considering the reliability, latency, and jitter of the evoked spikes^38,39^. Only neurons with reliable, consistent, and time-locked spikes upon light onset were considered tagged (see Methods). In total, 26 IT^Cux1^, 16 IT^PlxnD1^, 50 PT^Fezf2^, and 40 CT^Tle4^ neurons passed the quality criteria (**Fig. 4a-b**). For these tagged PNs, we observed reliable (*R* = 0.55 ± 0.21, mean ± SD) light evoked spiking with short latency (*L* = 4.9 ± 1.5 ms) and low jitter (*J* = 1.7 ± 0.6 ms) (**Fig. 4c, S6a-c**). Although each of these 4 PN types showed heterogeneity in activity, they exhibited overall distinct temporal patterns relative to RWD action sequence (**Fig. 4d-e**). Specifically, the activity of typical PT^Fezf2^ neurons rose before hand-lift, peaked during reach, and declined substantially by reach endpoint and during subsequent withdraw actions. Those of CT^Tle4^ typically ramped during the reach, peaked at reach-to-withdraw transition, and remained elevated during withdraw and drinking (**Fig. 4f-g**). Moreover, whereas PT^Fezf2^ activity was strongly tuned to the waterspout location, average CT^Tle4^ response showed only weak target location tuning (**Fig. 4h, S6d-e**). Overall, a large fraction of PT^Fezf2^ and CT^Tle4^ neurons correlated well with ongoing movement time series (**Fig. 4i**). As a comparison, ITs show a lower average discharge rate as compared with that of PT^Fezf2^ and CT^Tle4^ (**Fig. 4g**). These activity differences among PN types may explain the different behavior deficits obtained with inhibition during RWD. Notably, these results reveal different temporal activity patterns between the CT^Tle4^ corticothalamic and PT^Fezf2^ corticofugal output channels during RWD.

**Fig 4.**
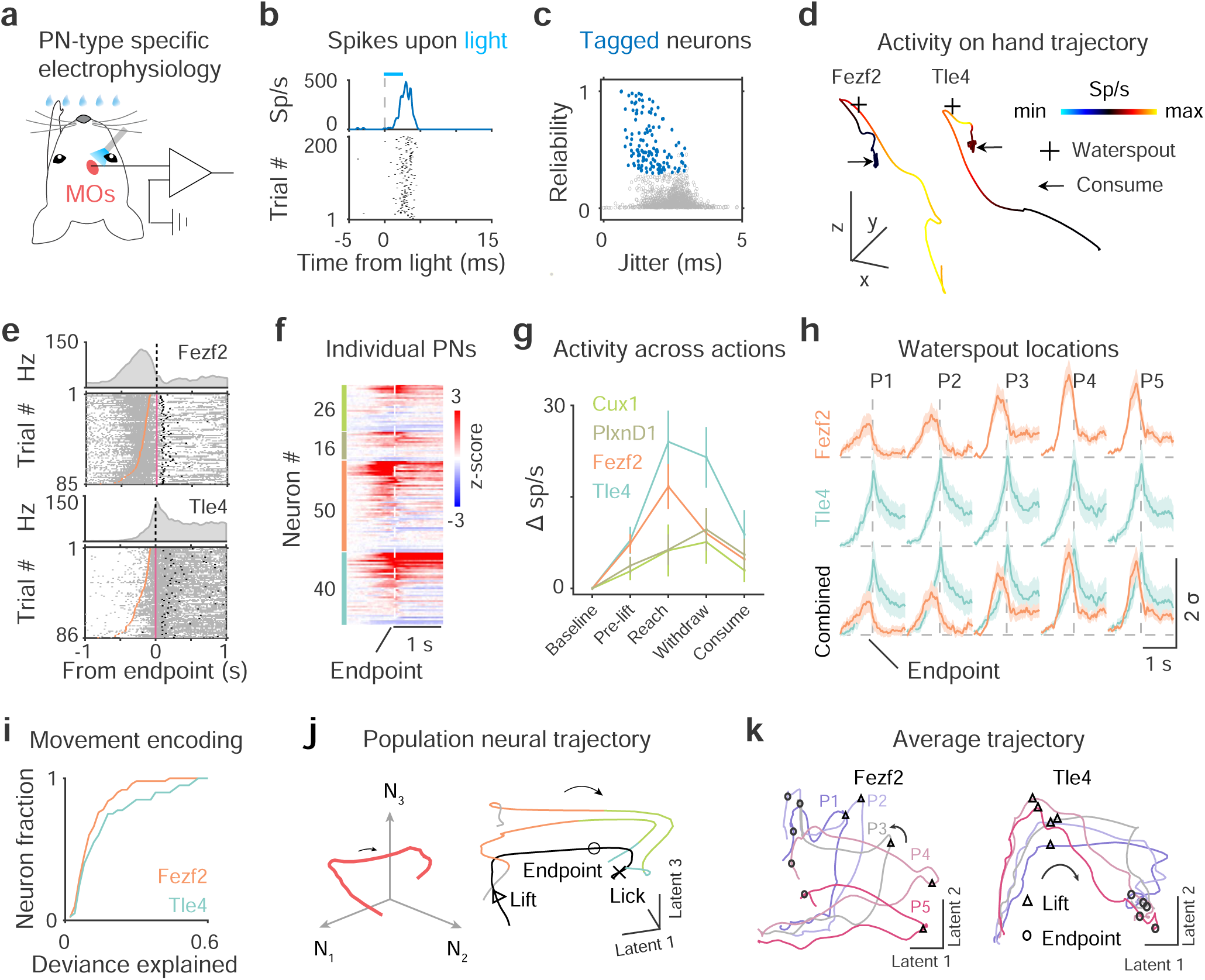
MOs-c PT and CT dynamics differentially correlate with action phase progression. **a.** Optogenetic tagging of MOs-c ChR2-expressing neurons with blue light pulses. **b.** Light-evoked peri-event time histograms (PETH, top) and raster (bottom) activity of a tagged neuron. Note the reliable and time-locked spikes relative to the onset of light pulses (dashed line) within 5 ms. Sp/s, spikes per second. **c.** Light-evoked spiking reliability and jitter of all neurons (gray circles). 132/1395 neurons were identified as tagged neurons (blue circles). **d.** Hand movement trajectory with neural activity (colormap) superimposed from a PT^Fezf2^ and a CT^Tle4^ neuron. x, forward; y, lateral; z, upward positions. Scale, 5 mm. **e.** Spike raster of tagged PNs during RWD. Trials are sorted by the duration between hand lift (orange ticks) and advance endpoint (pink ticks). Black ticks, first hand licks. **f.** Activity of tagged PNs aligned to advance endpoint (dashed line). Each row of the heatmap represents the z-score normalized activity of a neuron. Within a PN type, individual neurons are sorted by peak firing latency. (*n* = 26 IT^Cux1^, 16 IT^PlxnD1^, 50 PT^Fezf2^, and 40 CT^Tle4^) **g.** Baseline-subtracted activity across action phases of different PN types. The spikes during each phase were counted for each trial and averaged within a session for each PN. **h.** Average PN activity traces across five waterspout locations. Note the longer peak latency and more sustained firing in CT^Tle4^ than in PT^Fezf2^. **i.** Movement encoding performance as reflected by the deviance explained by Poisson-GLM models of individual neurons. (*n* = 50 PT^Fezf2^ and 40 CT^Tle4^.) **j.** Left, schematic showing state space population neural dynamics. Right, single-trial (color coded) and trial-averaged (black) neural trajectories of MOs-c of an example session in latent space. Colors indicate action phases of two example trials. **k.** Average PT^Fezf2^ and CT^Tle4^ population neural trajectories. P1-P5, five waterspout locations. Median lift, and advance endpoint time points are indicated by triangles and circles, respectively.

At the broad neural population level, simultaneously recorded MOs-c neural dynamics evolved with a smooth C-shaped trajectory with a clear “transitional bend” upon lift in a low-dimensional neural manifold along with the progression of RWD sequence (**Fig. 4j**). Among different trials, these trajectories shared similar geometric shapes but shifted in the latent space (low Procrustes distance, **Fig. 4j**). Interestingly, cell-type-targeted recording revealed that PT^Fezf2^ and CT^Tle4^ neurons also exhibited a smooth C-shape population trajectory with PT^Fezf2^ strongly modulated by target locations (**Fig. 4k**). On the other hand, IT^Cux1^ and IT^PlxnD1^ showed jerky population trajectories that varied according to target location (**Fig. S6f**). Given their categorically distinct axon projection patterns, these PN type-characteristic neural dynamics are likely separately conveyed to specific thalamic (CT^Tle4^) and corticofugal (PT^Fezf2^) target areas that contribute to ongoing movement control.

### MOs-c CT^Tle4^ enhances RWD-relevant thalamic dynamics

Our anatomical tracing revealed that MOs-c CT^Tle4^ axons terminate almost exclusively in the thalamus, which is distinct from pons-projecting PT neurons that project to multiple subcortical targets but with few thalamic collaterals (**Fig. 5a, S7a-b**). Specifically, MOs-c CT^Tle4^ neurons densely project to higher order motor thalamus including VAL and VM (**Fig. 5a**). In addition, retrograde mono-synaptic rabies tracing showed that MOs-c CT^Tle4^ neurons receive direct thalamic inputs from VAL, VM, PF, as well as long-range cortical inputs from MOp, SSp, SSs, ORBl, and subcortical inputs from GPe of the basal ganglia (**Fig. 5b, S7c-d**). Considering the anterograde projections of MOs-c CT^Tle4^ to VAL/VM (**Fig. 5a**), our results suggest a strong reciprocal loop between CT^Tle4^ and thalamic projection neurons (TPN). In addition to the reciprocal projection back to the MOs-c, TPN axons further send collaterals to MOp and SSp-ul in the RWD subnetwork as well as the striatum (**Fig. S7e**), consistent with results from single-cell reconstruction of TPNs^40^. Given that VAL/VM also receives inputs from the deep cerebellar nuclei, basal ganglia, and midbrain^41–46^ that convey ongoing movement state information, the “top-down” MOs-c CT^Tle4^ modulation likely regulates the integration of these inputs to TPNs and facilitate their output to influence the activity of a large network of cortical and striatal regions during RWD action sequence progression (**Fig. 5c**).

**Fig 5.**
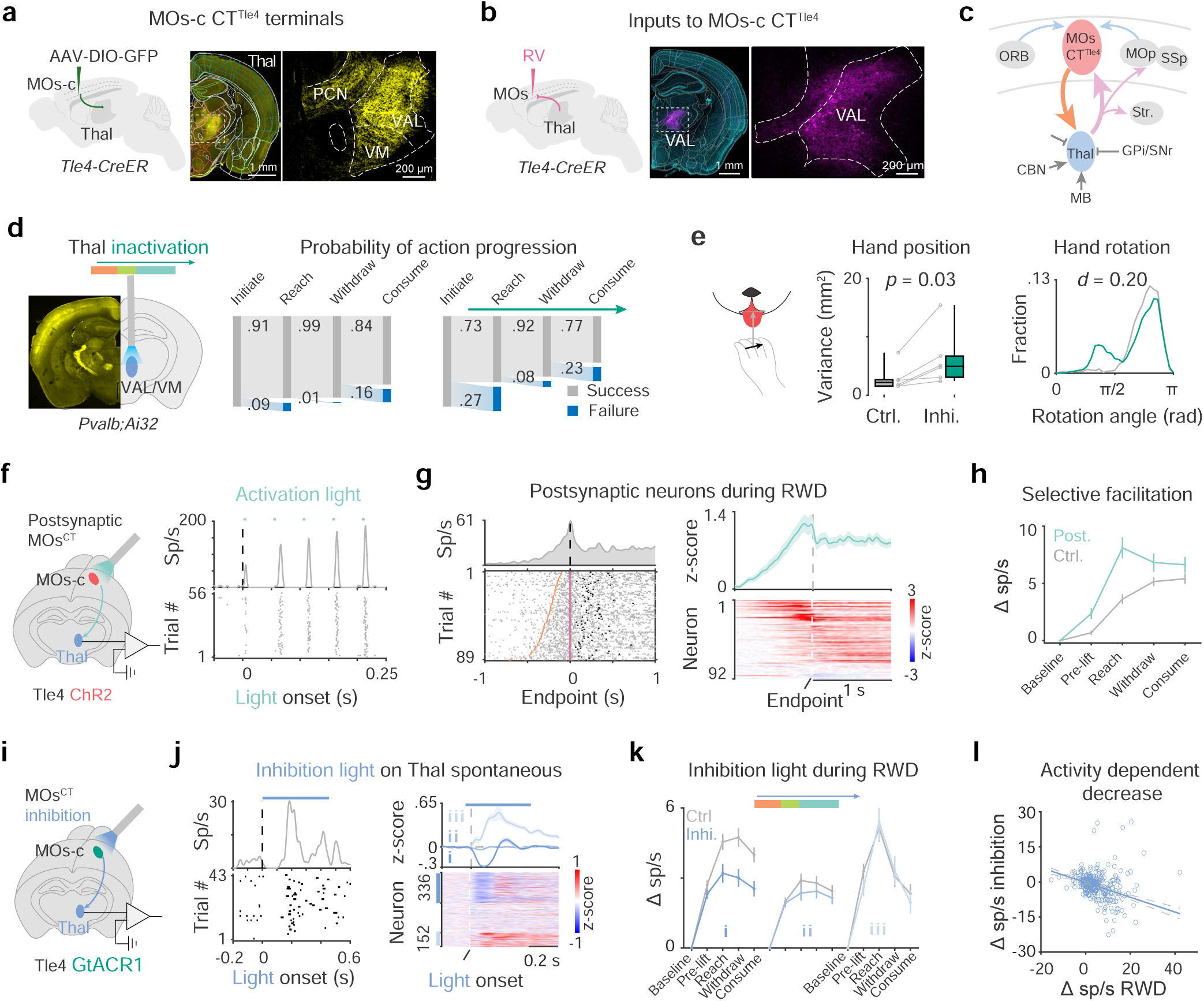
MOs-c CT^Tle4^ selectively enhances RWD-relevant thalamic dynamics. **a.** Neuron terminals of MOs-c CT^Tle4^ in higher-order thalamus. Right: zoom-in view of the boxed region (middle). VAL, ventral anterior-lateral complex; VM, ventralmedial thalamus; PCN, paracentral nucleus. **b.** Rabies (RV) tracing maps presynaptic inputs to MOs-c CT^Tle4^ from VAL. **c.** Schematics summary of the MOs-c corticothalamic reciprocal loop, which receives diverse cortical as well as subcortical inputs and projects to multiple cortical and striatal areas. Thal, thalamus; BG, basal ganglia, CBN, cerebellar nuclei; Str, striatum; OFC, orbitofrontal cortex. **d.** Interference of thalamic dynamics during reach impairs RWD action progression. Left, ChR2 expression (yellow) and optic fiber implantation into VAL/VM; Right, 15% (78/511) control trials and 42% (201/482) inhibition trials failed to complete the RWD sequence. (n = 6 sessions from 4 mice) **e.** Increased variance in hand position (left) and abnormal hand posture (right) upon lick with thalamic interference. (n = 6 sessions; Left: Wilcoxon rank sum test, \**p* < 0.05. Right: two-sample KS test, \*\*\**p* < 0.001.) **f.** Electrophysiological recording of MOs-c CT^Tle4^ postsynaptic neurons (TPN^Tle4-post^) in thalamus. Blue light pulses were applied to ChR2-expressing CT^Tle4^ neurons in the MOs-c (left). Light-evoked peri-event time histograms (top) and raster activity (bottom) of a TPN^Tle4-post^ (right). Note the facilitation of spiking with 20 Hz light pulses. **g.** RWD related activity of individual TPN^Tle4-post^. Left, spike raster of a TPN^Tle4-post^ during RWD. Right, summary of RWD related activity of 92 TPN^Tle4-post^. **h.** Increase in RWD-relevant activity of TPN^Tle4-post^ compared with control. (*n* = 92 TPN^Tle4-post^ and 210 control) **i.** Electrophysiological recording from thalamus upon optogenetic inhibition of MOs-c CT^Tle4^. **j.** Effect of MOs-c CT^Tle4^ inhibition (450 ms constant) on the spontaneous firing of thalamic neurons. Left: phasic decrease followed by brief increase of spontaneous discharge of a thalamus neuron during CT^Tle4^ inhibition. Right, neurons were divided into decreased (Group I, 336), non-modulated (Group II, 326), and increased (Group III, 152). **k.** Effect of MOs-c CT^Tle4^ inhibition on RWD-relevant thalamus activity among the three groups. Inhibition light was delivered in random half of trials around lift during RWD. **l.** Inhibition effect is dependent on RWD related increase in normal conditions. Each circle represents an individual Group I neuron. (*n* = 336 neurons, *R*^2^ = 0.25, *p* = 4.40×10^-22^)

Given the behavioral effects of MOs-c CT^Tle4^ inhibition, we hypothesized that normal VAL/VM activity is required during the RWD sequences. Thus, we perturbed thalamic activity by optogenetically activating GABAergic inhibitory terminals through an implanted optical fiber at the VAL/VM complex in *Pvalb;Ai32* mice (**Fig. 5d**). Similar to MOs-c CT^Tle4^ inhibition, closed-loop inhibition of VAL/VM activities upon lift significantly interfered with the progression of the RWD action sequence (**Fig. 5d**) and decreased hand-mouth coordination during consumption (**Fig. 5e**).

To explore how MOs-c CT^Tle4^ influences TPNs during RWD, we recorded spikes from VAL/VM complex and identified TPNs putatively postsynaptic to CT^Tle4^ (TPN^Tle4post^) by their consistent and time-locked spiking upon brief light activation of CT^Tle4^ (**Fig. 5f**). On average, light pulses evoked reliable (0.35 ± 0.19) spiking with a latency of 11.8 ± 2.4 ms (mean ± SD) and 3.8 ± 1.2 ms jitter in TPNs^Tle4post^ (92/647) (**Fig. S8a-c**). In response to a train of light pulses, the initial evoked spike frequency in TPNs^Tle4post^ was low but progressively facilitated with subsequent light pulses (**Fig. 5f, S8d**), consistent with the synaptic facilitation property observed in *in vitro* studies^47^. Such CT^Tle4^ TPN synaptic facilitation may dynamically modulate the spike output of TPN^Tle4post^ based on recent CT^Tle4^ spiking history^48^. While the activity of individual TPNs^Tle4post^ vary during RWD, the firing of many of them were tightly coupled to different action phases of RWD (**Fig. S8e**). Notably, TPN^Tle4pos^ discharges rose during the reach and were sustained during withdraw and drink, similar to those of MOs-c CT^Tle4^ (**Fig. 5g**). We observed a significant difference in RWD-related activity pattern between TPNs^Tle4post^ (92/647) and control TPNs (210/647) that were not modulated by MOs-c CT^Tle4^ (**Fig. 5h**). The RWD-related activity of TPNs^Tle4post^ was selectively amplified compared with control TPNs in an action-phase dependent manner (mixed-design ANOVA, *p* < 0.001, **Fig. 5h**). These results suggest that MOs-c CT^Tle4^ may selectively enhance the RWD-relevant output dynamics of a subset of TPNs.

Lastly, we tested whether CT^Tle4^ activity is required for the enhancement of thalamic dynamics during RWD. We optogenetically inhibited MOs-c CT^Tle4^ and recorded the neuron discharge in the VAL/VM complex (**Fig. 5i**). Inhibiting MOs-c CT^Tle4^ cells with a 450-ms light caused an initial decrease in the average spontaneous firing of thalamic neurons, followed by a brief increase above baseline activity, before returning to a level near the baseline (**Fig. 5j**), consistent with a previous observation^49^. Based on their differential response profiles following MOs-c CT^Tle4^ inhibition, we divided thalamic neurons into 3 groups: those that decreased activity (43% = 349/814, Group I), non-modulated (39% = 319/814, Group II) and those that increased activity (18% = 146/814, Group III) (**Fig. 5j**). We then examined the effect of MOs-c CT^Tle4^ inhibition on these 3 groups during RWD. Inhibition light was triggered around hand lift in half of the trials randomly. The light intensity was adjusted to a lower level (compared to those in **Fig. 3j-l)** so that a sufficient number of successful RWD trials were produced to compare inhibition vs control conditions. While mild MOs-c CT^Tle4^ inhibition resulted in an overall decrease in RWD-related thalamic activity (**Fig. S8f)**, individual TPNs among the 3 groups were differentially affected (**Fig. S8g**). In particular, Group I TPNs showed significant attenuation of discharge rate across RWD action phases (mixed-design ANOVA, *p* < 0.001, **Fig. 5k**), while the average firing frequency of the other TPNs (Group II and III) did not change compared with control trials (**Fig. 5k**). Notably, MOs-c CT^Tle4^ inhibition decreased Group I activity to a level comparable with that of Group II (mixed-design ANOVA, *p* > 0.05; **Fig. 5k**), suggesting that CT^Tle4^ inputs selectively enhance a subset of TPN activity during RWD. Moreover, Group I neurons with higher activity increase during RWD in control trials showed larger activity decrease upon CT^Tle4^ inhibition (**Fig. 5l**), suggesting that TPNs with higher RWD-relevant outputs are more dependent on CT^Tle4^ inputs.

As a subset of MOs-c PT^Fezf2^ also sends collaterals to the thalamus, we compared the effect of inhibiting these thalamus-innervating PT neurons (PT^Thal^) with that of inhibiting CT^Tle4^. We injected a *Flp*-dependent *AAV-retro-fDIO-Cre* in the thalamus of *Fezf2-Flp* mice followed by either a Cre-dependent AAV-DIO-mcherry or AAV-DIO-GtACR1 to specifically trace (**Fig. S9a-b**) or inhibit (**Fig. S9c)** PT^Thal^. The fractions of decreased (11% = 92/849), non-modulated (52% = 439/849) and increased (37% = 318/849) thalamic neuron groups were significantly different from that of CT^Tle4^ inhibition (Chi-squared test, *p* < 0.001, **Fig. S9c**). Overall, MOs-c PT^Thal^ inhibition resulted in a delayed increase of TPN average spontaneous activity (**Fig. S9c)**. PT^Thal^ inhibition during RWD did not significantly decrease the RWD-relevant activity of thalamic neurons (**Fig. S9d**). It is possible that the additional dense MOs-c PT^Thal^ innervation of the basal ganglia and zona incerta (ZI) nucleus (**Fig. S9b**), which provide inhibitory signals to the thalamus, makes the *in vivo* effect of PT^Thal^ on TPNs different from that of CT^Tle4^. Altogether, these results highlight the unique role of CT^Tle4^ in facilitating the RWD-relevant thalamic dynamics.

## DISCUSSION

The mouse reach-withdraw-drink behavior is representative of the many forelimb-mediated skillful consummatory behaviors of rodents and primates in that it entails the coordinated progression from the allocentric reach-grasp to egocentric withdraw-drink to achieve an ethological goal^1^. Using cell-type genetic tools for wide-field imaging and optogenetic inhibition, we identify a key high-order area MOs-c, embedded within a dynamic cortical network, that orchestrates the progression and coordination of RWD actions. Cell-type-targeted recording and manipulation within the MOs-c reveal that pyramidal tract and corticothalamic output channels show distinct activity dynamics during RWD and differentially contribute to action sequence progression and coordination, with a particularly prominent and unexpected role of the corticothalamic pathway. Notably, MOs-c CT^Tle4^ manifest sustained dynamics across RWD action phases and selectively enhance the RGD-relevant activity dynamics of their postsynaptic thalamus neurons, which also contribute to action progression and coordination. MOs-c CT^Tle4^ receive converging inputs from forelimb and orofacial sensorimotor areas of the RWD network and are reciprocally connected to their thalamic neurons, which project back to this cortical network. Therefore, we discover the crucial role of a corticothalamic loop, which may selectively amplify the thalamic integration of diverse cortical and subcortical sensorimotor streams to promote action progression and coordination in skilled motor behaviors.

### MOs-c is crucial in the progression and coordination of RWD actions

Previous studies have identified various primary and higher order motor areas in the planning, execution, and sequencing of forelimb^2,16,17,50,51^ and orofacial^18,20,21,52,53^ movements. Here, using an unbiased PN-type resolution survey of cortex-wide activity patterns during a complex motor sequence, we identified multiple cortical areas within a dynamic sensorimotor network that correlate with the progression of RWD behavior. Among these areas, inhibition of MOs-c resulted in deficits not only in the progression of forelimb actions but also the coordination between forelimb and mouth movements, suggesting a major role in the articulation of RWD. MOs-c partially overlaps with the rostral forelimb area (RFA), a broad premotor region in rats^54^ and mice^35,50^. This area is densely connected with both forelimb and orofacial sensory and motor areas^36,55,56^, making it well-poised to monitor and orchestrate the RWD action sequence. We suggest that MOs-c might be analogous to primate premotor and/or supplemental motor areas implicated in sensory-guided coordination of complex movement^14,57–59^. Beyond RWD, MOs-c might function more broadly to orchestrate cross-body action coordination in other complex behaviors. In addition to MOs-c, SSp-ul inhibition also resulted in significant deficits in RWD, suggesting a significant role of sensory processing in this behavior. We note, however, that MOp inhibition resulted in more subtle deficits, suggesting a lesser role of primary forelimb motor area in this behavior. How MOs-c communicates with other network nodes (e.g. MOp, SSp-ul) and with subcortical structures during RWD is of major interest for future investigations.

### MOs-c output channels and the crucial role of the corticothalamic pathway

MOs-c communicates with other cortical and striatal regions through intratelencephalic (IT) neurons and broadcasts subcortical output through extratelencephalic (ET) neurons^23^. Our imaging, electrophysiology, and optogenetic analyses of multiple major PN projection types consistently pinpoint a more significant contribution of ET than IT subpopulations to RWD, and further identified the distinct roles of two cell-type-specific ET channels. Most previous studies of cortical output pathways in motor control have been directed toward L5 PT neurons^60,61^. Indeed, L5 PT have been variously implicated in the planning, initiation and execution of movement^50,62–65^. Notably, L5 PT extend collaterals to subsets of high-order thalamic regions and strongly innervates subpopulations of TPNs^24,40,64^. Based on anatomical and mostly *in vitro* physiology evidence, these PT collaterals are thought to represent a “driver” type input that mediates the transthalamic (cortico-thalamo-cortical) pathways implicated in sensory processing^28,30,66,67^. In contrast, L6 CT dominate the corticothalamic inputs by covering the entire thalamus and innervating most if not all TPNs^68^. They are considered “modulator” inputs based on their smaller and weaker synapses directed to distal dendrites of TPNs^26,69^. A recent study suggests that CT neurons in the primary motor cortex exert a permissive role in motor execution, likely through intracortical feedforward inhibition of PT neurons^70^. Here we have discovered a novel function of the L6 corticothalamic pathway in high-order motor cortex in regulating action progression and coordination in a complex motor sequence. Whereas PT^Fezf2^ activity rises during reach then declines substantially during withdraw, CT^Tle4^ activity rises sharply during reach and grasp then remains elevated throughout withdraw and the subsequent hand-lick events. Consistent with this pattern, whereas PT^Fezf2^ inhibition perturbs reach and grasp, CT^Tle4^ inhibition additionally interferes withdraw and hand lick. These results implicate PT^Fezf2^ in the targeting and progression of forelimb actions, especially when reaching for more difficult locations. Importantly, they highlight a crucial role of CT^Tle4^ in the entire sequence of reach, grasp, and withdraw including their coordination with the oral actions to consume the water.

PTs^Fezf2^ likely exert their function through regulating their numerous subcortical target structures^50,64,71^, thus a mechanistic understanding would require a systematic dissection of their projection diversity as well as disentangling the role of each target. In contrast, the exclusive subcortical target of CT^Tle4^ unambiguously suggests a role for the motor thalamus in action progression and coordination. We note that CT^Tle4^ also extends local axon collaterals^22^ that may impact cortical circuits^72,73^, a subject of future investigation.

### Cortico-thalamo-cortical communication in action progression and coordination

The high-order motor thalamus VAL/VM receives major afferents from the basal ganglia, cerebellar nuclei, and midbrain structures in addition to top-down inputs from cortical motor areas^41–46^. They thus can integrate subcortical movement-related motivational, body state, and sensory information with cortical descending streams that convey motor plan and feedback signals^74^. The outcome of this “super-integration” is then conveyed by TPN spike trains to the recurrent as well as divergent cortical and striatal networks^29,75^. Previous studies implicate high-order thalamus in the planning^76,77^, initiation^78^, and execution^27,79^ of movements. In particular, ongoing thalamic activity is necessary for driving cortical dynamics during forelimb reaching^27,77^ and singing^80^. Here we show that perturbing VAL/VM activity interferes with RWD action progression and hand-mouth coordination. Consistent with in vitro findings^47^, signaling from MOs-c CT^Tle4^ to their postsynaptic TPNs (TPN^Tle4post^) involves short-term facilitation. Notably, the activity dynamics of VAL/VM TPNs^Tle4post^ showed significantly larger activity rise during reach that sustains during withdraw and drink as compared with TPNs not modulated by MOs-c CT^Tle4^. The action-phase selective enhancement of the RWD-relevant dynamics in TPN is dependent on ongoing MOs-c CT^Tle4^ dynamics, as closed-loop inhibition of MOs-c CT^Tle4^ prevented this enhancement. Together, these results suggest that ongoing MOs-c CT^Tle4^ dynamics is required for amplifying action-phase relevant activities of a subset of TPNs, whose outputs may further influence the RWD-related cortical network dynamics that facilitate action sequence progression and coordination.

Our rabies tracing reveals that MOs-c CT^Tle4^ neurons receive converging inputs from several sensorimotor areas of the RWD subnetwork and are reciprocally connected with VAL/VM thalamic target neurons, which in turn project back to cortical areas within and beyond the RWD subnetwork. Considering that VAL/VM receive rich subcortical inputs from the basal ganglia, cerebellar nuclei, and midbrain ^41–46^, the CT^MOs-c^−TPN^VM/VAL^ reciprocal loop is embedded within the larger cortical-basal ganglia/cerebellar-thalamic loop systems. In this context, a prominent feature of both MOs-c CT^Tle4^ and VAL/VM TPNs^Tle4post^ activity dynamics is their sustained firing across the RWD action sequence; their reciprocal excitatory connectivity and the CT^Tle4^-TPNs^Tle4post^ short-term synaptic facilitation may promote this property. It is possible that the persistent activity dynamics in the MOs-c corticothalamic loop may be primed or driven by successive rounds of action-related cortical and subcortical inputs while powerfully modulated by concurrent top-down CT^Tle4^ activity. With output to multiple sensorimotor cortical and striatal areas, this corticothalamic loop activity pattern may facilitate the temporal evolution of system-wide neural dynamics that underlie action progression and coordination during reach-withdraw-drink.

## ACKNOWLEDGEMENTS

We thank Drs. Nuo Li, Matthew Kaufman, Court Alan Hull, Stephen Lisberger for comments on the manuscript; Tatiana Engel, Yanliang Shi, William Galbavy, Katherine S. Matho, and Dhananjay Huilgol for discussions on the project; Anne Churchland and Simon Musall for advice on building wide-field imaging setup; Priscilla Wu, Joshua Hatfield, and Baoxia Han for animal preparation and maintenance; Weixin Zhong for schematic drawings. This work was supported by the NIMH research project U19MH114823-01 grant to Z.J.H. Z.J.H. is supported by an NIH Director’s Pioneer Award 1DP1MH129954-01.

## AUTHOR CONTRIBUTIONS

Y.L. and Z.J.H. conceived the study. Z.J.H. acquired the funding, managed the project, and supervised the study. Y.L. designed experiments, built setups, performed experiments, established pipelines, analyzed the data, and presented them for visualization. X.A. shared resources and collected RV-tracing data. P.J.M trained animals, collected brains, and analyzed electrophysiological data. Y.Q. prepared AAV vectors and helped with electrophysiological recording. X.H.X contributed to behavioral training and optogenetic manipulation. S.Z. packaged AAVs and performed *in situ* hybridization. H.M. helped with the wide-field imaging rig. S.M.S interpreted anatomical data. L.B.R. and N.B. analyzed and interpreted electrophysiological data. I.Q.W. contributed to behavioral annotations and analysis. Z.J.H. and Y.L. wrote the manuscript with significant edits from I.Q.W. All authors discussed the data, interpreted the data, offered advice for data analysis, reviewed, and provided inputs to the manuscript.

## DECLARATION OF INTERESTS

The authors declare no competing interests.

## INCLUSION AND DIVERSITY

We support inclusive, diverse, and equitable conduct of research.

## RESOURCE AVAILABILITY

### Animals and materials availability

All mice, reagents and materials are openly available with detailed source information listed in the Methods.

### Data and code availability

All original data reported in this study and any additional information required to reanalyze the data reported in this paper will be shared upon request. Customized code related to data collection and analysis would be are available on Zenodo (https://zenodo.org/records/10041374; https://zenodo.org/records/10041296).

## METHODS

### Mice

Animal care, use, surgical and behavioral procedures conformed to the guidelines of the National Institutes of Health. The experiments were approved by the Institutional Animal Care and Use Committee of Cold Spring Harbor Laboratory and Duke University. Experiments were conducted with 8-week-to 16-week-old male and female mice. The number of animals used in each experiment is noted in the corresponding section. Mouse strains were: *Emx1-Cre* (JAX#005628), *Cux1-CreER* (JAX#036300), *PlxnD1-CreER* (JAX#036294)*, Fezf2-CreER* (JAX#036296), *Fezf2-Flp* (JAX#036297), *Tle4-CreER* (JAX#036298), *Ai148D* (JAX#030328), *Pvalb-IRES-Cre* (JAX#017320), *Ai32* (JAX#024109). Mice were housed in groups of up to five mice per cage, in a room with a 12/12 light/dark cycle. After surgery, mice were housed in a new home cage individually or with familiar groups for at least one week prior to the experiments.

### Surgery

Materials, including instruments used in surgery, were sterilized, and stereotaxic surgery was performed using aseptic techniques. Surgical anesthesia was maintained using 1%-2% isoflurane via inhalation. The analgesic drug ketoprofen (5 mg/kg, subcutaneous) effective for up to 24 hours, was administered prior to the beginning of the surgery. The local anesthetic lidocaine was administered subcutaneously at the intended incision site (2-4 mg/kg). Body temperature was maintained at 37° C using a feedback-controlled heating blanket. Eye ointment was applied to prevent the eyes from drying. After disinfecting with betadine solution (5-10%) and ethanol (70%), a small incision of the scalp was created to expose the skull.

A titanium flat headpost was implanted for head restrained experiments. For wide field calcium imaging, the skull was cleaned with saline, and a thin layer of cyanoacrylate glue (Zap-A-Gap CA+, Pacer Technology) was applied on the skull to clear the bone. After the cyanoacrylate glue cured, cortical blood vessels were clearly visible. Then, a circular flat head post was attached to the skull using dental cement (C&B Metabond, Parkell; Ortho-Jet, Lang Dental) leaving most of the dorsal cortex exposed. For inhibition screening experiments, a thin skull preparation^81^ was used in *Pvalb-IRES-Cre*;*Ai32* mice. Clear low toxicity silicone adhesive (KWIK-SIL, World Precision Instruments) was applied on the dorsal cortex as protection against dust and scratches. Animals were allowed to recover from surgery before the experiments began.

Viral vector injections were performed using a Nanoliter 2010 injector (World Precision Instruments) controlled by a SMARTouch controller at a rate of 46 nL/min. Cortical injections were targeted to MOs-c (AP +1.6 mm, ML 1.4 mm). For thalamic injections, pipette tip was targeted to VAL/VM (AP-1.1 mm, ML1.1 mm, DV 3.7 mm). Virus includes: *AAV_DJ_-CAG-DIO-GtACR1-EYFP* (2.43×10^13^ vg/mL, Vigene Biosciences); *AAV-EF1a-DIO-hChR2-EYFP* (2.3×10^13^ vg/mL, Addgene#35509); *AAV-retro-EF1a-Flp* (2.3×10^13^ vg/mL, Addgene#55637); *AAV-retro-fDIO-Cre* (2.3×10^13^ vg/mL, Addgene#121675); *AAV9-CAG-DIO-EGFP* (2.5×10^13^ vg/mL, Addgene#51502); *AAV9-EF1a-fDIO-mCherry* (2.3×10^13^ vg/mL, Addgene#114471). 300-500 nL volume of virus was used. After injection, the glass pipette was left in place for ten minutes and then slowly withdrawn at a speed of 50 μm per min. Optical fibers (200 μm, 0.37 NA; RWD Life Science Inc.) fitted into an LC-sized ceramic fiber ferrule were implanted. After 12 to 21 days of waiting time for post-surgery recovery and viral expression, the animals were used in experiments.

For retrograde monosynaptic rabies tracing from MOs-c, we first injected the starter virus of *AAV8-hSyn-FLEX-TVA-P2A-eGFP-2A-oG* (400 nL, >3.64×10^13^ vg/mL, Addgene#85225) into MOs-c. Three weeks later, the same mice were injected in the MOs-c with *EnVA-dG-Rabies-mCherry* (500 nL, >1.0×10^8^ vg/mL, Addgene#32636, Salk GT3 Vector Core). Brain tissue was prepared for histologic examination 7-10 days after the rabies virus injection.

### Tamoxifen induction

The temporally controlled expression of Cre recombinase was achieved by intraperitoneal injection of tamoxifen (two 100 mg/kg injection at 20 mg/ml, prepared in corn oil) of *CreER* knock-in driver lines. For *Ai148D* reporters crossed with *CreER* mice, the first induction was on the day of weaning and the second induction was one week later. For Cre recombinase induction in virus injected *CreER* mice, two injections of tamoxifen were administered intraperitoneally on day 1 and day 3 after virus injection (day 0).

### Head-restrained reach and withdraw to consume

The day before behavioral training, animals were weighed and moved to a new cage with new bedding and food but with restricted access to water. Animals received supplemental water to meet daily water needs to maintain body weight >80% of the initial weight, monitored by daily weighing and evaluation.

The reach for water task^31^ was controlled in real-time with customized MATLAB (MathWorks) code. A data acquisition board (USB-6351; National Instruments) was used to communicate between the software and hardware (piezo sensors, water valves, and linear actuators). Two high-speed USB cameras (FL3-U3-13S2C-CS or BFS-U3-04S2C-CS; FLIR) acquired video data from the front and left side of the mouse. The cameras were synchronized and calibrated to enable three-dimensional infrared recording of the animal’s forelimb and orofacial movements. Simultaneous acquisition and storage of the video at 240 frames per second (fps) at a resolution of 640×480 pixels were achieved using a customized Bonsai workflow. Water delivery information for each trial and touch sensor data were shared in real-time between the MATLAB control code, the cameras, and electrophysiological recording systems.

Mice were trained to reach for water in two phases, within which they were required to obtain a hit rate > 80%. In phase 1, the waterspout was fixed on the left side of the animal’s snout, and in phase 2, it was moved to one of 5 equidistant locations (identified as left P1, P2, center P3, right P4, P5) with P3 centered and each location approximately 3 mm apart in front of the animal’s nose. Phase 1 training consisted of 1 session each day for 3 days, with 100 trials per session. Pretraining to reach involved placing a waterspout (made from a 21-gauge needle and providing a drop of sucrose solution, 10% w/v, 20∼50 μL) ∼3 mm to the left side of the snout midline. The waterspout tip was horizontally aligned with the upper point of the animal’s mouth, which is about 4 mm below the tip of the nose. Animals were trained to use their left hand to grasp the water drop with their right limb blocked. The waterspout tip was initially close to the mouth but then gradually moved away to 3-5 mm from the animal’s mouth. Water was delivered at a random inter-trial interval (12-20 s). The random duration was long enough for the animal to replace its hand to the starting position after consuming water on each trial. The start position of the hand was 20-30 mm posterior and downward from the waterspout tip. A piezo sensor detected waterspout contact events. If the animal failed to reach within 8 seconds after water delivery, a new trial began after a 30 second timeout. For phase 2 training, a linear actuator (L16-R Miniature Linear Servo for RC; Actuonix Motion Devices) moved the waterspout to one of the 5 locations.

### Cortex-wide calcium imaging

All mouse driver lines were bred with reporter strains for calcium imaging or electrophysiological recording except for *Emx1-Cre* mice. They were injected retro-orbitally with *AAV-PHP.eB-CAG-DIO-GCaMP7f* (2.46×10^13^ vg/mL, Addgene) at postnatal days 14 because they failed to breed with *Ai148D* mice.

Cortex-wide calcium imaging^33,82,83^ was performed with an inverted tandem-lens macroscope in combination with a scientific complementary metal-oxide semiconductor (sCMOS) camera (Edge 5.5, PCO) with a wide field. The focal length of the top lens was 105 mm (DC-Nikkor, Nikon) and that of the bottom lens was 85 mm (85M-S, Rokinon), resulting in a magnification of ×1.24. The total field of view was 12.4 mm by 10.5 mm with a spatial resolution of ∼20 μm/pixel. To capture GCaMP fluorescence, a 525 nm band-pass filter (#86–963, Edmund optics) was placed in front of the camera. Using alternating excitation light at two different wavelengths, calcium-dependent fluorescence was isolated and corrected for intrinsic signals (for example, hemodynamic responses). Excitation light was projected onto the cortical surface using a 495 nm long-pass dichroic mirror (T495lpxr, Chroma) placed between the two macro lenses. The excitation light was generated by a collimated blue LED (470 nm, M470L3, Thorlabs) and a collimated violet LED (405 nm, M405L3, Thorlabs) that were coupled into the same excitation path using a dichroic mirror (#87–063, Edmund optics). The alternating illumination between the two LEDs and the acquisition by the imaging camera were controlled by an Arduino Uno R3. The camera ran at 50 fps, producing one set of frames with blue excitation and another set with violet excitation, each at 25 fps. The exposure state of each frame was recorded. Excitation of GCaMP at 405 nm resulted in non-calcium-dependent fluorescence, allowing isolation of the true calcium-dependent signal by subtracting fluorescence changes in violet frames from the blue illumination frames by regression, as detailed below. Subsequent analyses were based on this differential signal at 25 fps.

### Optogenetic manipulation of behavior

The open-source guide was used to achieve real-time and closed-loop control based on markerless hand position tracking^84^. Briefly, real-time reach behavior was monitored using a USB camera (Flea3; Point Grey) on the left side of the mouse, ipsilateral to the reaching forelimb. A trained deep neural network with ResNet-50 model (the same one used for behavior analysis) for the side view video was embedded in a custom Bonsai workflow to trigger optogenetic stimulation based on real-time detection of hand position (https://github.com/bonsai-rx/deeplabcut). Low-latency control of light was achieved with videos capturing at 25 fps and a resolution of 640 × 480 pixels on a Windows workstation equipped with a GeForce RTX 2080 Ti GPU (NVIDIA). A 200 μm optical fiber delivered 473 nm blue (SSL-473-0100-10TM-D, Sanctity Laser) or 532 nm green (SSL-532-0200-10TM-D, Sanctity Laser) light.

Inactivation of different cortical areas or thalamus was achieved with *Pvalb-IRES-Cre*;*Ai32* mice that allow optogenetic activation of local PV interneurons for an inhibition screening^85^. Blue light pulses (5 ms, 50 Hz, 473 nm) were triggered in 50% of reach trials, as the animal’s real-time hand position crossed a predefined threshold in a closed-loop manner. The light spot size was restricted with a 200 μm optical fiber, with its tip directly contacting the thinned skull. The fiber ferrule was positioned with an MP-285 micromanipulator (Sutter Instrument). Light intensity at the tip was adjusted to 5 mW. The triggered light was automatically turned off four seconds after the water delivery.

For neuron-type specific inhibition experiments, the inhibitory opsin GtACR1 was locally expressed with AAV and illuminated with green light (532 nm) adjusted to 10 mW. Two optogenetic inhibition strategies were applied. 1) Closed-loop reach photoinhibition: the light was on when the hand moved across a predefined position during reaching and off when the hand repositioned at the start location. 2) Prolonged inhibition: light was turned on 1 second before water delivery and lasted for the entire trial. Experiments involved either bilateral or unilateral inhibition.

### Multielectrode array recording

The surgery procedure is as described in previous sections. To provide a ground reference, a mini screw connected to a silver wire (A-M systems) was implanted into the skull above the left visual cortex. Before the first recording session, a craniotomy was performed under isoflurane anesthesia. A linear silicon probe was slowly lowered into the cortex with an MP-285 micromanipulator (Sutter Instrument). A thin layer of clear silicone elastomer (Kwik-Sil, World Precision Instruments) or agarose was applied over the craniotomy after the electrode was positioned to the desired position to stabilize the exposed brain. The brain was allowed to settle for 15-30 minutes before recordings began. At the end of the recording session, the probe was retracted, and the craniotomy was sealed with Kwik-Sil to allow a subsequent session on the following day.

Extracellular spikes were recorded using linear 32-channel silicon probes (ASSY-37 H4, Cambridge NeuroTech, or A1×32-5mm-25-177, A4×8-5mm-100-200-177, NeuroNexus) or Neuropixels 1.0/2.0 probes *in vivo* combined with optogenetic stimulation of ChR2-expressing neurons. For optogenetic tagging of ChR2-expressing neurons in the cortex, probes were inserted to the cortex, and stimulation light was applied to the cortex locally. To identify postsynaptic neurons in the thalamus, probes were inserted to the recipient thalamus while the stimulation light was applied to the ChR2-expressing cortex through the cleared skull. For 32-channel probes, voltage signals were continuously recorded at 32 kHz by a Digital Lynx 4SX recording system (Neuralynx). Raw data was collected and saved for analysis using Cheetah software. Neuronal activity was band-pass filtered (300-6000 Hz) for real-time visualization of optogenetic light evoked effect for optical tagging. For Neuropixels, data were acquired via PXIe-1083 acquisition box (National Instruments) with open-source software SpikeGLX (http://billkarsh.github.io/SpikeGLX) sampled at approximately 30 kHz calibrated for each probe. To localize probe tracks, probes were coated with fluorescent dye CM-Dil (1 μg/μL in ethanol; C7000, Thermo Fisher) before each recording. The fluorescent tracks of probes were imaged and further registered to Allen Institute Mouse Brain CCF coordinate system with the SHARP-Track (https://github.com/cortex-lab/allenCCF). Neuron depth information was estimated with the registered coordinates.

### Signal synchronization

Systems for behavioral recording, cortex-wide calcium imaging, and electrophysiological recordings were synchronized with a common synchronization signal.

### RNA *in situ* and immunohistochemistry

After the experiments, animals were euthanized with isoflurane and perfused with saline followed by 4% paraformaldehyde fixation using a peristaltic pump. Brains of *PlxnD1-CreER;Ai148* mice were dissected, fixed, and cut into 10 µm sections. For RNA *in situ* hybridization chain reaction, brain sections were hybridized with *Fezf2* probes (Molecular Instruments) in probe hybridization buffer at 37 °C for 24 hours, washed with probe wash buffer, and incubated with an amplification buffer at 25 °C for 24 hours in a 24-well plate. For immunohistochemistry after RNA *in situ*, the same brain sections were stained with rabbit anti-Cux1 (1:500, 11733-1-AP, Proteintech) or mouse anti-Tle4 antibody (1:500, Cat#sc-365406, Santa Cruz Biotechnology). Briefly, brain sections were first pretreated with 10% Blocking One (Cat#03953-95, Nacalai Tesque) in PBS with 0.3% Triton X-100 (Blocking solution) at room temperature for 1 hour, then incubated with primary antibody in Blocking solution at 4 °C overnight. The brain sections were washed with PBS the next day and incubated with Cy3 conjugated donkey anti-rabbit (1:500, Cat#711-165-152, Jackson Immunoresearch) or donkey anti-mouse (1:500, Cat#715-165-150, Jackson Immunoresearch) secondary antibody for 2 hours at room temperature. Brain slices were then mounted on slides for confocal imaging (ZEISS Axio Observer).

### Data processing and analysis

Data processing and analyses were performed with MATLAB (Mathworks) or Python, unless otherwise specified. Sufficient reach-to-grasp trials were collected for each condition such that all results could be reproduced robustly. No statistical methods were used to predetermine sample size.

#### Extraction of movement time series

The reaching behavior was monitored by two high-speed cameras (Flea 3, Teledyne) at 240 fps from both the front and the left side of the animal. The geometric relationship between the two cameras was calculated using the Camera Calibrator app in MATLAB. Real-time videos of the behavior session were acquired and saved using a customized code in Bonsai (version 2.6) for offline analysis. Two deep neural networks were trained separately using DeepLabCut 2.0^32^ (https://github.com/DeepLabCut) to track the body part positions in front and side views. Neural network training was performed using over 2000 frames (1040 front and 1076 side frames) from 20 different recording sessions of 20 individual animals. Images of different animal colors, body sizes, head-restrained setups, illumination conditions from different behavior phases were included to train a relatively robust network. In total, 18 keypoints in the front view and 22 keypoints in the side view were labeled on each frame. These included the digits, nose, mouth, tongue, waterspout, and the water drop. For the front view, 1,030,000 training iterations were achieved with a training error of 2.07 pixels and test error of 3.89 pixels at a statistical confidence level of > 0.95. For the side view, 1,030,000 training iterations produced a training error of 1.74 pixels and test error of 4.16 pixels at a confidence level of > 0.95. The 3D positions of the hand and waterspout were reconstructed through stereo triangulation. Only samples with a network predicting confidence level of > 0.95 were used for analyses. In cases of missing samples, the corresponding samples from a cubic spline were used to fill the trajectory (gaps > 100 ms were not used).

#### Quantification of action phases

The RWD behavior is a closed loop act in which mice monitor the target location online using olfactory and tactile cues to direct their reach^31,86^. The movement of reach, grasp and withdraw and their constituent actions were identified using a previously described movement classification scheme^87^. The reach consists of lift, aim and advance segments and directs the hand to the waterspout. The grasp occurs with the end of advance and consists of opening and extending the digits and then closing them to purchase a water drop. The withdraw is an egocentric movement in which the hand is supinated immediately after grasp and further supinated as the hand retracts to the mouth for drinking. The following constituent action segments and critical events (e.g. start and end points of an action) of the RWD are featured in the automated kinematic analysis pipeline for pose estimation after keypoint extraction by DeepLabCut.

##### Reach

was quantified as successful lift, aim and advance of hand towards target.

###### Lift

The hand was raised from the resting position and partially pronated with the digits collected (i.e. lightly closed and flexed). The 3D reconstructed position of digit 3 was used to represent and track hand movement. The first frame in which the vertical hand speed increased above 75 mm/s (upward) defined lift initiation. Left-hand speed was the absolute value of the derivative of 3D left-hand position. The lift phase consisted of the time series from lift initiation to peak speed.

###### Aim

The hand was positioned by an elbow-in movement of the upper arm, the digit pointing direction was rotated and the palm aimed toward the waterspout. Hand/digit rotation was characterized using the rotation vector that connects the midpoint and tip of digit 3 and digit 4 in the front view (**Fig. S1b**). The vector connecting moment-by-moment hand position and the waterspout position was defined as the range vector (vector ***r*** in **Fig. S1c**). The moment-by-moment direction and amplitude change of range vector reflected the hand movement in relation to the waterspout. The waterspout aiming angular deviation (angle *δ* in **Fig. S1c**) was the angle between the current finger pointing direction (vector ***p*** in **Fig. S1c**) and the instantaneous range vector direction at a given moment. A waterspout aiming score was defined as the cosine value of the angular deviation (cos(*δ*) in **Fig. S1c**). Aim completion was defined as the time point when waterspout aiming score is higher than 0.866 (δ < π/6). The aiming phase was defined as the time points from first hand peak speed point to aim completion.

###### Advance

The hand was advanced toward the waterspout by upper arm movement and opening of the elbow with concurrent opening and extension of digits. The advance phase was defined as the time from aim completion to advance endpoint which terminates just before grasp. If an animal failed to reach the waterspout or to grasp water, the hand would usually return to the aim position with the digits closed and flexed for another advance.

###### Advance endpoint

The vector direction from the initial hand position to the waterspout was the reference direction (dashed line in **Fig. 1f**). The median hand position during the 2 seconds before water delivery (pre-lift position) was used as the initial hand position. Real-time hand to spout distance was calculated as the projection of the range vector onto this reference direction. The farthest reaching point along the reference direction defined the reaching endpoint. At this point the hand was positioned adjacent to the waterspout with the digits extended and the palm in a near vertical orientation in preparation for grasping. The length of the rotation vector (**Fig. S1d**) reflects how much the digits were abducted and is defined as digit-open size. The amplitude and direction of the range vector at the reaching endpoint were characterized to represent reaching endpoint accuracy.

##### Withdraw

Withdraw-to-consume phase was the time points between the advance endpoint and the first hand-lick. *Grasp* involves the closing of digits to purchase a water drop after contacting the waterspout. Grasp completion was defined as the time point when tips of all four digits become invisible (confidence level < 0.5) from the lateral view following hand-open and digit-extended state. The waterspout contact event was identified by the piezo sensor attached to the waterspout. The hand was then withdrawn to the mouth by upper arm movement that lowered the hand to the level of the mouth with the palm in a vertical orientation. It was further *supinated* by movement at the wrist such that the palm faces upward. Supination was measured as wrist rotation to a position in which the palm faces up with the hand rotation score higher than 0.5 (*θ* < π/3). The rotation score was the cosine value of the direction of the line connecting midpoints of digit 4 and digit 3 in the front view relative to horizontal direction (cos(*θ*) in **Fig. S1f**).

##### Consume

Animals consumed water by licking their left hands. Tongue protrusion events were identified by the trained deep neural network for front view videos with confidence level > 0.95. Most tongue protrusions occurred after animals successfully grasped the waterspout. The hand was positioned near the mouth and made repositioning movements that included digit opening and extending in coordination with licking. The median position of the left and right mouth corners in that session was the mouth position. The distance between moment-by-moment hand position and the reference mouth position in the front view was used as hand to mouth distance. The distance between the hand and tongue (***d*** in **Fig. S1e**) and the hand rotation score (***s*** in **Fig. S1f**) upon tongue protrusion quantified the coordination between hand and tongue movement for drinking.

##### Other orofacial movements

Prior to reaching, animals detected the water by sniffing and orienting their nose toward the corresponding waterspout location. Moment-to-moment nose displacement movements were obtained by comparing the median value of nose positions with the left side displacement being positive and right displacement being negative. Mouth open and close movements were identified by calculating the area covered by two mouth corners and upper lip in the front view video.

##### Replace

After drinking, the hand reversed movement direction, lowered from reaching or from the mouth, and returned to the approximate starting position.

#### Cortex-wide movement encoding model

Methods to extract and decompose cortex-wide calcium dynamics were described previously^82,83,88^. The landmarks of the dorsal cortex were marked, and the mask was set in the scope of the dorsal cortex from an example frame. Next, control and GCaMP frames were split from raw videos. Each cropped video frame (size 440 × 440) was transformed to a flat array. Images from different trials were then concatenated resulting in a two-dimensional matrix (size *n* × *t*, *n* = 440 × 440, *t* is time frame). Denoising was performed with an SVD-based method (singular value decomposition). SVD returned ‘spatial components’ *U* (of size pixels by components), ‘temporal components’ *V*^T^ (of size components by frames) and singular values *S* (of size components by components). To reduce computational costs, all subsequent analyses were performed on the product *SV*^T^ represented as *V_c_*. Results of the analyses on *SV*^T^ were later multiplied with *U* to recover results back to the original pixel space. The denoising step outputs a low-rank decomposition of *Y_raw_* = *UV_c_* + *E* represented as an *n* × *t* matrix; here *UV_c_* is a low-rank representation of the signal in *Y_raw_*, and residual *E* is considered noise. The output matrices *U* and *V_c_* are much smaller than the raw data *Y_raw_*, leading to compression rates above 95%, with minimal loss of the visible signal. Finally, an established regression-based correction method isolated a purely calcium dependent signal by subtracting the control channel signal *Y_v_* (405 nm illumination) from the GCaMP channel signal *Y_g_* (473nm illumination). With the corrected values for each trial, the signal values in a time window one second before water delivery were used as the baseline to calculate z-score. All wide field imaging data was registered to the Allen reference mouse brain Common Coordinate Framework (CCF3) using five anatomical landmarks: the left, center and right points where anterior cortex meets the olfactory bulbs, the medial point at the base of retrosplenial cortex, and Bregma, labeled manually for each imaging session during mask setting.

Generalized linear encoding models (GLM) with ridge regularization were built to predict the cortex-wide neural dynamics with moment-by-moment animal behavior (behavior matrix *M*, size frames by 13)^82^. Cortex-wide neural dynamics were represented by all ‘temporal components’ *V_c_* (size components by frames) saved after SVD decomposition. The model was fitted using ridge regression with 10-fold cross-validation to avoid overfitting. The regularization penalty was estimated separately for each component of *V_c_* data on the first fold of validation and used the same value for other folds of validation. A newly modeled variable *V_m_* of the same size as *V_c_* was predicted using this GLM process and used to compute Y*_m_* = *UV_m_* as predicted pixel-wise neural dynamics. The predicted *Y_m_* was compared with *Y_raw_* and pixel-wise explained variance (*R^2^*) was obtained to quantify the cross-validated GLM performance (represented with cv*R*^2^).

To construct the behavioral matrix *M*, movement time series related to hand, digit, wrist, and orofacial movements were derived from the frame-by-frame labeled points in front and side videos. In addition to the isolated left-hand movements, the hand relationships to the waterspout and to the mouth were included to fully capture the events and their spatiotemporal relationships that constitute RWD. Thirteen analog behavioral variables after kinematic analysis were selected to describe hand, digit, wrist, and orofacial movement during RWD. Those ethologically meaningful behavioral variables were not necessarily orthogonal to each other. Normalized behavior variables were down-sampled to match cortex wide activity, combined to make matrix *M* (size frames by 13), and used to predict cortex-wide neural dynamics. Thirteen continuous variables are listed as follows: 1) forward position of left hand, 2) upward position of left hand, 3) lateral position of left hand, 4) path length moved from left hand onset, 5) moving speed of left hand, 6) digit open size of left hand, 7) moving speed of right hand, 8) nose displacement, 9) mouth open size, 10) supination score of left hand, 11) left hand to mouth distance, 12) waterspout aiming score by digits of left hand, and 13) left hand to waterspout distance. The first 6 variables represent the kinematics of left-hand movement. Variables from 7 to 9 depict the movement of other body parts. The last 4 variables (from 10 to 13) reflect the relationship between the left hand and target or between the left reaching hand and mouth. Reaching forelimb-related variables (1-6 and 10-13) were used to predict calcium dynamics. We also tried predicting neural activity with all thirteen movement variables. Similar results were observed. No time-shifted versions of movement time series were tested.

#### Cortical nodes

Masks for regions of interest (ROIs) were derived by thresholding averaged calcium activity amplitude and the GLM encoding model performance (cv*R*^2^). All neuron types were considered. Activity amplitude or GLM performance larger than 80% of the maximum value of all pixels on the dorsal cortex was used as the high threshold. Activity amplitude lower than -0.4 z-score was used as the low threshold. ROI 1 (MOs-c, central region of secondary motor cortex, centered at AP/ML: +1.60/±1.37 mm) and ROI 5 (Prt, parietal cortex including part of SSp-ll and SSp-tr, -1.18/±1.63 mm) were overlapping regions with activity amplitude larger than the 80% of maximum activity from *Emx1*, *Fezf2* and *Tle4* populations. ROI 4 (SSp-ul, anterior-lateral forelimb somatosensory cortex and the unassigned region, +0.23/±2.62 mm) was observed in *Tle4* activity larger than high threshold and further isolated with GLM performance of *Emx1*, *Fezf2* and *Tle4* populations. ROI 3 (MO-orf, orofacial motor cortex in the lateral part of the anterior cortex, +1.67/±2.08 mm) covered the pixels that were lower than -0.4 z-score in *Cux1* activity. ROI 6 (SSp-bfd, anterior part of barrel field and nose primary somatosensory cortex, -0.71/±2.69 mm) switched hemisphere as the waterspout was moved from ipsilateral to contralateral side of the animal from both *Cux1* and *PlxnD1* populations. ROI 2 (MOp-ul, primary motor cortex for upper limb, +0.41/±1.69 mm) was revealed by the extended *Fezf2* activity into MOp on the right hemisphere as compared to the left hemisphere. Because the border of some ROIs overlapped, the coordinates of the center of each ROI relative to Bregma were shown above and used to guide further experiments. The median value of pixels within 100 μm diameter from the center of a ROI was used to represent its activity.

#### Waterspout location selectivity

Waterspout location selectivity reflects the difference in firing rate when animals reach for differently located waterspout. Location selectivity (LS) was calculated as *LS* = (*r*_ipsi_ – *r*_contra_) /*r*_avg_; where *r*_ipsi_ is the firing rate during reaching for ipsilateral targets relative to the reaching hand; *r*_contra_ is the firing rate during reaching for contralateral targets; *r*_avg_ is the average across all waterspout locations. For wide-field imaging data, pixel-wise *LS* values were provided at specific behavioral time points. For electrophysiological data, *LS* values of individual neurons were shown across different action phases.

#### Spike sorting, quality control and optogenetic tagging

Saved raw electrophysiological data was rearranged by the channel depth, median-subtracted across channels and time, and the results were saved in 16-bit binary files for further spike detection and sorting using Kilosort2 or Kilosort4 software^89^ (https://github.com/MouseLand/Kilosort). A proper probe configuration was created using default parameters for spike detection and sorting. Spikes were further visualized and manually curated in phy2 (https://github.com/cortex-lab/phy) to remove apparent noise. Sorted data was analyzed using custom MATLAB scripts. Besides spike shape, several metrics were taken into consideration for cluster quality control^90^: median amplitude (> 60 μV), refractory period violation rate (< 15%), and noise cutoff. In optogenetic tagging experiments, neurons were further curated by only keeping those with correlation coefficient of spike waveform >= 0.85 between random selected spontaneous spikes and light stimulation evoked spikes. Only ‘good’ clusters with mean spike firing frequency over 0.5 Hz were used for further analysis.

Optogenetic tagging was adopted to confirm the cell type of recorded neurons with methods described previously^38,39,91^. During the first and last five minutes of the recording session, when animals were not engaged in the behavioral task, 473 nm blue light pulses (2 ms or 5 ms duration) at different frequencies (0.1, 10 and 20 Hz) were delivered through an optical fiber over the craniotomy for optical tagging. Light intensity was adjusted based on the amplitude of light evoked response during the process. To determine whether a neuron was activated by light, a statistical *p* value was computed by comparing the distribution latencies of first light evoked spikes within 8 ms from light onset and a permuted latency distribution of spontaneous spikes. To further characterize the tagged neurons, we computed additional metrics including the ***reliability*** of light-evoked spiking (*R*), median ***latency*** of triggered spikes after light onset (*L*), and the standard deviation of the latency across different light pulses as ***jitter*** (*J*) for each neuron. Only ‘good’ neurons with *p* < 0.05 and *R* >= 0.3 were considered as optogenetic tagged neurons. The spike latency and reliability of ChR2-expressing neurons upon light onset depends on many factors including the stimulation light intensity, opsin expression level, the cellular compactization of illumination, the kinetics of light sensitive opsin, electrical properties and connections of recorded neurons.

Similar methods were used to determine whether a postsynaptic neuron was activated by cortical projection neurons. As the process involves synaptic transmission, the statistical *p* value was computed by searching for the first spike within a 20 ms window from light onset (*p* < 0.05). Only ‘good’ neurons with *p* < 0.05 and *R* >= 0.1 were considered as postsynaptic neurons. Only *p* < 0.05 without R >= 0.1 restriction resulted in similar conclusions.

#### PETH

To compute peri-event histograms (PETH) aligned to different behavioral events, the spikes of individual neurons were binned in 10 ms windows and smoothed with a Gaussian kernel (20 ms standard deviation). Average spike rates across trials were used to represent the activity of individual neurons. All spike rates were z-score normalized to the mean and standard deviation of a pooled distribution of binned baseline activity (1 second before water delivery) across all trials. Each unit had twenty PETHs related to four behavioral events: hand lift onset, advance endpoint, first lick, and last lick, each with five different target locations.

Non-parametric Wilcoxon signed-rank tests were used for each neuron to determine whether a neuron is significantly increased, decreased, or not modulated. Activity change during 600 ms around the advance endpoint (300 ms before and 300 ms after advance endpoint) was compared to baseline.

#### Spike-based movement decoding and encoding models

To decode movement time series, the spiking activity of simultaneously recorded neurons from each session was used to predict thirteen behavioral variables contained in the behavioral matrix *M* individually with Gaussian GLM (*glmfit*) within a session. Spiking activity was binned and smoothed with a 20 ms window for each neuron. Behavioral profiles were down sampled to match spiking activity. No lagged versions of movement time series were tested. Either normalized activity or the principal components explaining over 85% of the variance after PCA in each session was used as regressors, which resulted in the same conclusion.

To reveal the encoding information by individual neurons, we used Poisson GLM (*glmfit*) to predict the spike count within 20 ms bins with movement time series (behavior matrix *M*)^92^.

To avoid overfitting, similar procedures were used to perform cross-validation for both decoding and encoding models, in which 25% of samples were left out in a random trial-based manner with trained GLM on the remaining data. The trained model was used on the left-out sample to predict movement profiles. The predicted movement profile was compared with original behavioral variables to compute the proportion of explained deviance/variance in representation of the model performance. We repeated this procedure 100 times and averaged the explained deviance/variance as cross-validated GLM performance for the session. The proportion of deviance explained in Poisson models is a similar parameter as the variance explained in a Gaussian model (*R*^2^) to measure model performance.

Naive Bayes classifiers were used to predict either five waterspout locations directly or three waterspout location classes relative to body side^93^. For decoding across behavioral phases, spiking activity during action phases from individual trials within a session was used.

We used CEBRA to decode nonlinearly embedded action phases in population neural activity^94^. Spikes of simultaneously recorded neurons that passed quality control were binned at 1/240 s. Each bin was annotated with different action phases. The four behavioral windows for each trial are as follows: 1) pre-lift, the 200 ms window before lift onset; 2) reach, the window from lift to advance endpoint; 3) withdraw-to-consume, the window from advance endpoint to the first consumption hand lick; 4) consume, the window from the first consumption lick to the last consumption lick in the trial. Time points from different reach trials were concatenated. We used a nonparametric supervised learning k-nearest neighbor algorithm as a decoding method for CEBRA. We also mapped the projection of simultaneously recorded spikes onto the movement time series with CEBRA and superimposed different time points with action phase information, which also showed clear embedding of progression of actions.

#### Population neural trajectory

Concatenation was used to visualize and quantify time-evolution of trial-averaged population neural activity for different waterspout locations^17,27,95^. Briefly, a data matrix of neural activity *X* size *n* × *ct* was compiled where *n* was the number of neurons, *c* was the number of waterspout locations, and *t* was the number of time points (at 20 ms bins). This matrix contained the firing rates of every neuron for every condition and every analyzed time point. Principal component analysis (PCA) was used to decompose the centered and normalized *X* from *n* × *ct* to *m* × *ct* with *m* orthogonal dimensions with MATLAB function *pca*. The normalized *X* were projected to the *m* orthogonal dimensions to get *score* (*m* × *ct*) values for all time points along all dimensions. To visualize the time-evolution of population activity, we plotted the projection of the normalized *X* along the first three principal components.

To track the evolution of neural dynamics through time for each trial, we projected the responses of simultaneously recorded neurons of individual trails into a low-dimensional space with Gaussian-process factor analysis (GPFA) ^96^. The dissimilarity in geometric shapes between a pair of neural trajectories was computed using the best shape-preserving Euclidean transformation between the two trajectories using the MATLAB *procrustes* function. The Procrustes distance was quantified for each pair of neural trajectories at different waterspout locations. The lower values of Procrustes distance reflected higher similarity in shapes.

## SUPPLEMENTAL INFORMATION

**Supplementary video 1**

RWD behavior, 3D-reconstructed hand trajectories, and selected movement time series across different waterspout locations are shown. Movement components, including reach (lift, aim, advance), withdraw and drink, are superimposed on hand trajectory. Movement time series include hand to target distance, moving speed, rotation score and waterspout aiming score of the left reaching hand. Example trials from three waterspout locations were arranged sequentially. The movie is played at 1/40 of the original behavior speed.

**Supplementary table 1**

Detailed statistical information.

---

**Fig S1.**
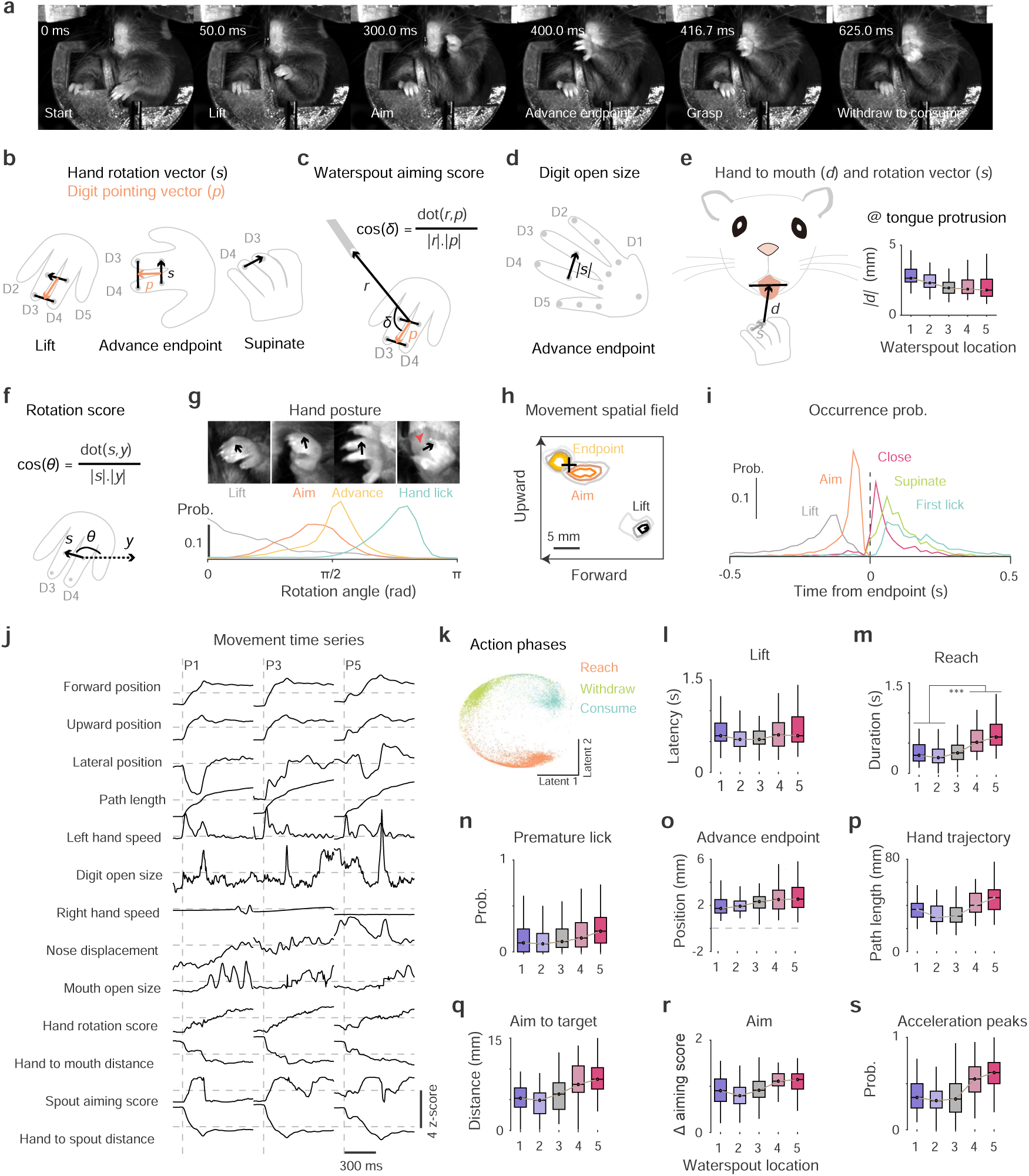
Characterization and quantification of RWD behavior. Related to Fig 1. **a.** Video frames (front view) showing hand locations in a representative trial at different time points during the reach. **b.** Schematic of the measurement of hand rotation direction (black vector) and finger pointing direction (orange vector) as represented by key points on left digits. The hand supinates to a horizontal position with the lift, advance and to a palm up position for licking. **c.** Quantification of waterspout aiming score (cos(*δ*)) to measure the hand posture for aim location and posture toward waterspout. **d.** Quantification of digit open/extend size for waterspout grasp, quantified as the length of the hand rotation vector. **e.** Schematic for quantifying hand to mouth distance (*d*) and hand rotation vector (*s*) upon lick. The hand was maintained close to the mouth and supinated during tongue protrusions. **f.** Schematic for quantifying the hand rotation score (cos(*θ*)) relative to the horizontal reference vector (*y*). Value 1 means the palm faces up and is fully supinated. -1 indicates the palm faces down and fully pronated. 0 means left hand is vertical and facing towards the right. Typically, the hand rotation score is close to -1 before lift, 0 at grasp and close to 1 at hand lick. **g.** Top: hand posture progression at lift (the fingers slightly closed and flexed), at aim (palm rotated toward target), at advance endpoint (fingers extend and open for grasp) and at hand-lick (hand can be closed or open). Bottom: distribution of hand rotation angle as a reflection of palm-facing direction. Data from 6229 lifts, aims, advance endpoints; and 119584 licks from 70 sessions in 25 mice. Note the near 180-degree hand supination from reach-grasp (pronated) to withdraw-lick (supinated). **h.** Spatial contour map of hand locations (side view) at lift, aim and advance endpoint. Contours indicate probability starting from 0.01/mm^2^ with equal increment of 0.01/mm^2^; 6229 trials from 70 sessions in 25 mice across five target locations; +, waterspout location. **i.** Occurrence probability of sequential RWD movements relative to the reach endpoint. Results of 3924 trials from 70 sessions in 25 mice reaching for P2. **j.** Hand and oral movement time series for waterspout positions P1, P3 and P5. **k.** Mapping of the action phases with movement time series. Each dot is a movement time point in the latent space. **l–s.** Behavior modulation by waterspout locations: latency to lift (**l**), reach duration (**m**), premature lick before waterspout contact (**n**), reach end position relative to waterspout (**o**), path length (**p**), distance from aim onset of reach to the target (**q**), change of digit aiming score during aim (**r**), probability of significant acceleration peaks during aim and advance (**s**). 70 sessions from 25 mice. Median ± interquartile range. Horizontal lines in boxplots indicate 75%, 50%, and 25% percentile. Whiskers represent data point span to 90% or 10% percentile. See Supplementary table for statistics.

**Fig S2.**
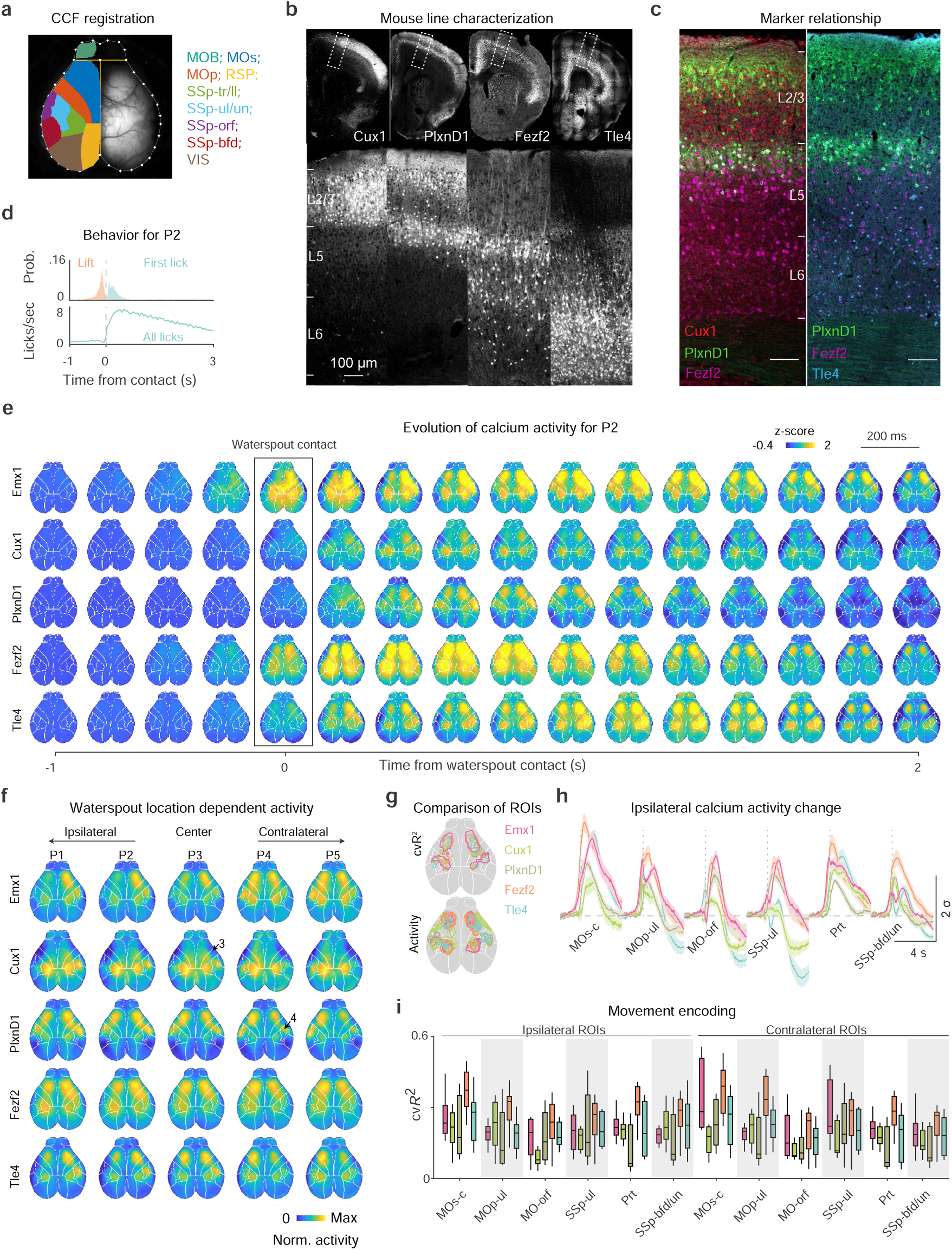
Cortex-wide calcium activity during RWD. Related to Fig 2. **a.** Registered atlas areas superimposed on an image of the dorsal cortex. MOB, main olfactory bulb; MOs, secondary motor cortex; MOp, primary motor cortex; RSP, retrosplenial cortex; SSp, primary somatosensory cortex; tr, trunk; ll, lower limb; ul/un, upper limb and unknown region; orf, orofacial; bfd, barrel field; VIS, visual cortex. White dots outline the cortex; yellow horizontal line shows olfactory bulb and neocortex boundary; yellow vertical line shows the midsagittal suture.ds. **b.** Coronal brain sections showing the laminar pattern of PN types labeled by driver lines crossed to reporter mice. Bottom row is zoom-in, and rotated view of the dashed box annotated with layers L1-L6. **c.** PN markers across cortical layers. Immunostaining of Cux1 (red) and Tle4 (blue), was carried out after mRNA in situ hybridization of *Fezf2* (magenta) in *PlxnD1;Ai148* (green) mice. Scale bar, 100 μm. **d.** Temporal occurrence of RWD movements after training at location P2. Top, lift and first hand-lick probability (Prob.) distribution relative to waterspout contact. Bottom, average lick frequency aligned to waterspout contact. (*n* = 3924 trials from 70 sessions in 25 mice) **e.** Average sequential activity frames at 200 ms intervals centered on waterspout contact (black box) during RWD from P2. (*n* = 9 sessions from 5 mice for PN^Emx1^; 7 sessions from 4 mice for IT^Cux1^; 11 sessions from 4 mice for IT^PlxnD1^; 12 sessions from 6 mice for PT^Fezf2^; 10 sessions from 5 mice for CT^Tle4^.) **f.** Average cortex-wide calcium activity at the five target locations during the whole RWD process. Note the increase of ipsilateral (left) hemisphere activity as the target moved from P1 to P5. **g.** Summary and comparison of ROIs. Top: ROIs by thresholding of normalized GLM performance. Bottom, ROIs by thresholding of normalized activity. **h.** Calcium activity change of ipsilateral ROIs. **i.** Quantification of the performance of the GLM encoding model for different ROIs.

**Fig S3.**
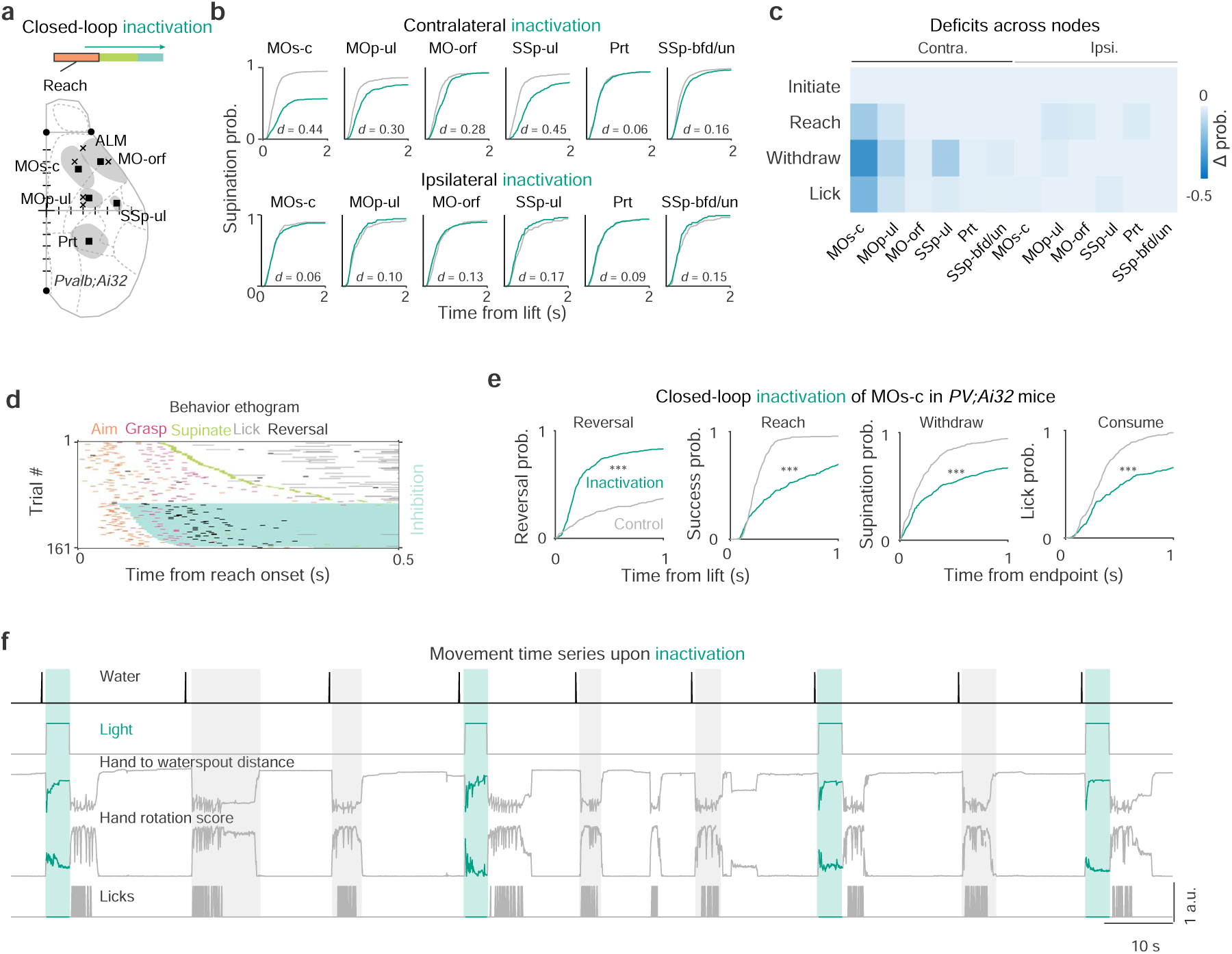
Photoinhibition survey across cortical nodes during RWD. Related to Fig 3. **a.** ChR2-assisted closed-loop photoinhibition of cortical areas during RWD. Crosses (‘x’) represent the center of several previously characterized cortical areas, from anterior to posterior: anterior lateral motor cortex (ALM; Guo, 2014), rostral forelimb area (RFA; Tennant, 2011), rostral forelimb orofacial area (RFO; An, 2023), caudal forelimb area (CFA; Tennant, 2011), and primary motor cortex for the upper limb (Sauerbrei, 2020; Muñoz-Castañeda, 2022). Black squares indicate the centers of region of interest in this study. +, bregma; scale, 0.5 mm. **b.** Cumulative distribution of supination latency relative to hand lift in closed-loop inhibition and control trials of each node in contralateral and ipsilateral hemisphere. *d*, two-sample Kolmogorov-Smirnov distance. (*n* = 5 mice, see Supplementary table for statistics) **c.** Changes in occurrence probability of constituent movements between inhibition and control trials for each contra- and ipsi-lateral cortical node. Performance of component movements, reach, withdraw, and drink were quantified with the probability of successful waterspout contact, full hand supination, and lick respectively. (*n* = 5 mice) **d.** Closed-loop inhibition of contralateral MOs-c impaired action progression represented as an ethogram from an example session. Actions accomplishment is color coded. The relative onset and duration of the inhibition light for each trial are indicated with cyan shades. Note inhibition attenuates the completion of the RWD sequence. **e.** Closed-loop MOs-c inhibition resulted in increased hand reversals during reach, decreased target contact after lift, supination after grasp, and hand lick after grasp. (*n* = 5 mice, two-sample KS test.) **f.** Movement time series during closed-loop MOs-c inhibition. Exemplary hand movements in relation to the target, hand rotation, and lick from several consecutive control (gray) and inhibition (tortoise) trials are shown. Movement profiles are normalized. Note that upon termination of inhibition, animals immediately resumed and completed the action sequence of RWD.

**Fig S4.**
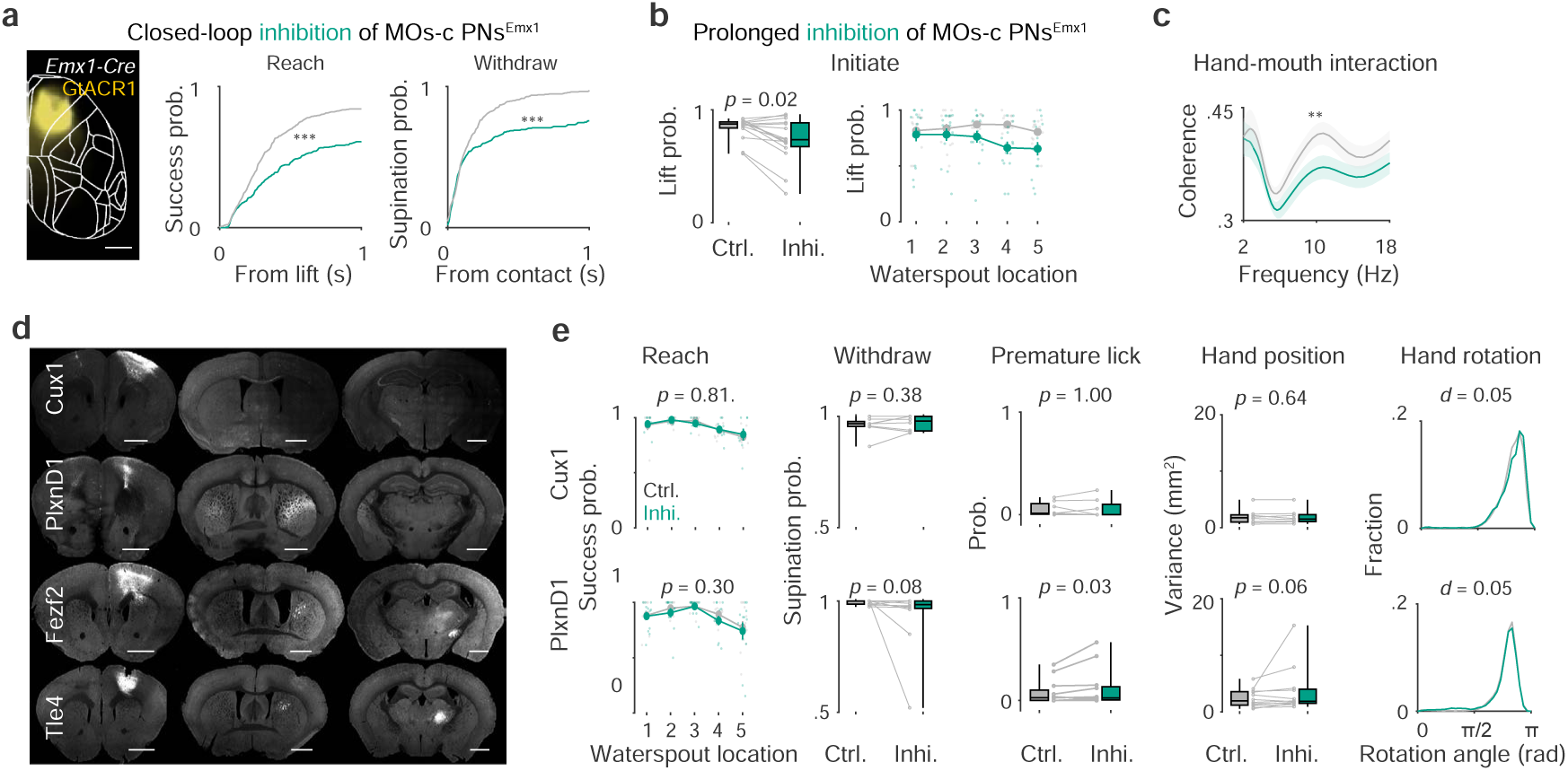
MOs-c PN-type-specific contribution to RWD. Related to Fig 3. **a.** Left, viral expression of the inhibitory opsin GtACR1 in MOs-c of an *Emx1-Cre* mouse. Scale, 1 mm. Right, impaired reach and withdraw upon closed-loop inhibition of MOs-c PNs. (*n* = 5 PN^Emx1^ mice.) **b.** Decrease of lift probability to initiate reaching for contralateral targets upon prolonged MOs-c PN inhibition (*n* = 15 sessions from 8 PN^Emx1^ mice. Left, Wilcoxon rank sum test. Right, mixed-design ANOVA, inhibition *F*_1,56_ = 10.74, *p* < 0.01; inhibition × target *F*_4,56_ = 4.46, *p* < 0.01). **c.** Decreased coordination between hand upward-downward movement and mouth open-close movement during drinking measured by their coherence (Mixed-design ANOVA, inhibition *F*_1,494_ = 12.96, \*\**p* < 0.01). **d.** Cortical, striatal, and thalamic projections from different MOs-c PN types. IT neurons show projection to the contralateral cortex and no projection to thalamus. Note the rare projection of IT^Cux1^ to striatum as compared with that of IT^PlxnD1^. PT^Fezf2^ and CT^Tle4^ both project to thalamus. Scale, 1 mm. **e.** Effect on reach, withdraw, and consumption upon prolonged IT inhibition (*n* = 8 sessions from 6 IT^Cux1^ mice and 12 sessions from 7 IT^PlxnD1^ mice. Reach: mixed-design ANOVA; IT^Cux1^ *F*_1,28_ = 0.06, *p* = 0.81; IT^PlxnD1^ *F*_1,44_ = 1.2, *p* = 0.30. Boxplots for supination probability after waterspout contact, premature lick and variance of hand position upon licks: Wilcoxon rank sum test. Plots for hand rotation upon licks: two-sample KS test)

**Fig S5.**
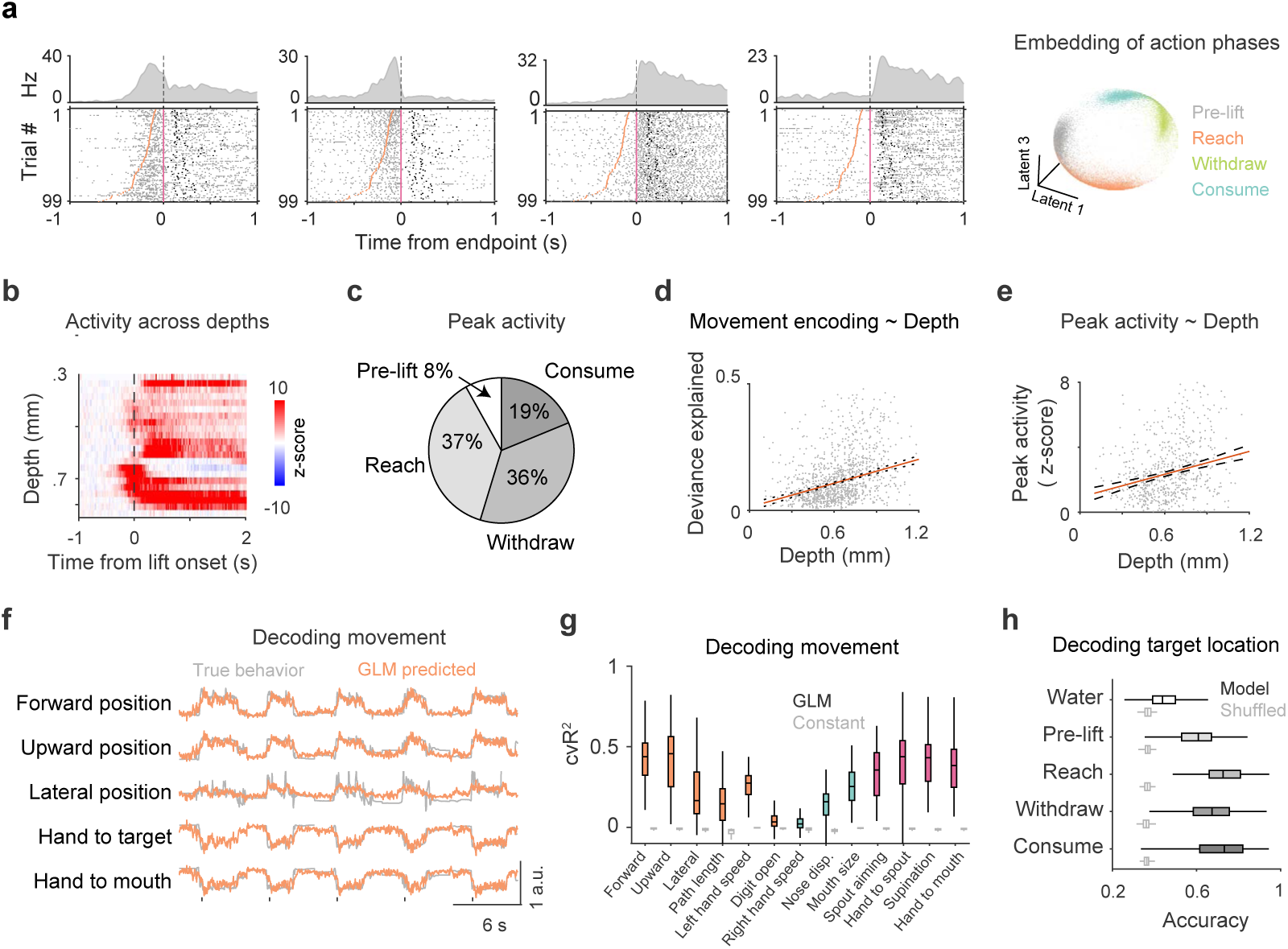
Electrophysiological recording from MOs-c during RWD. Related to Fig 4. **a.** Action phase encoding in simultaneously recorded individual and population neurons in MOs-c. Raster panels show spikes of individual neurons in relation to actions. Right most panel shows action phase embedding in the latent space of population neural activity revealed by supervised CEBRA model in an example session (*n* = 108 neurons). Each dot represents a time point (1/240 s). Superimposed colors indicate different action phases. **b.** Neuronal activity at different cortical depths relative to hand lift (0) from an exemplar session. **c.** Fraction of activated neurons that peaked at different action phases during RWD. **d.** Increase of movement encoding as the neural depth increases. *R*^2^ = 0.157, *p* = 9.73×10^-44^. Movement encoding of individual neurons was represented by the deviance explained by a Poisson-GLM. See Methods. **e.** Increase of peak activity as neural depth increases. *p* = 1.48×10^-20^. **f.** Correlation between exemplar actual movement time series (gray) and GLM-predicted traces (orange) using simultaneously recorded spiking activity from an exemplar recording session. Movement time series include forward, upward, lateral hand position, and hand position in relation to waterspout or mouth. Spikes were binned in 20 ms bins to predict time series using GLM. **g.** Population MOs-c activity decodes forelimb movement times series. Note the higher correlation with forelimb movement kinematics (orange) in relation to waterspout or mouth (magenta), but lower correlation with movements of other body parts (blue). Gray boxplots indicate cross-validated performance of the null model as control. (*n* = 106 sessions) **h.** MOs-c activity decodes target locations indicated with cross-validated decoding accuracy of Naïve Bayes classifiers. Unfilled box plots indicate performance of the shuffled model as control. (*n* = 106 sessions)

**Fig S6.**
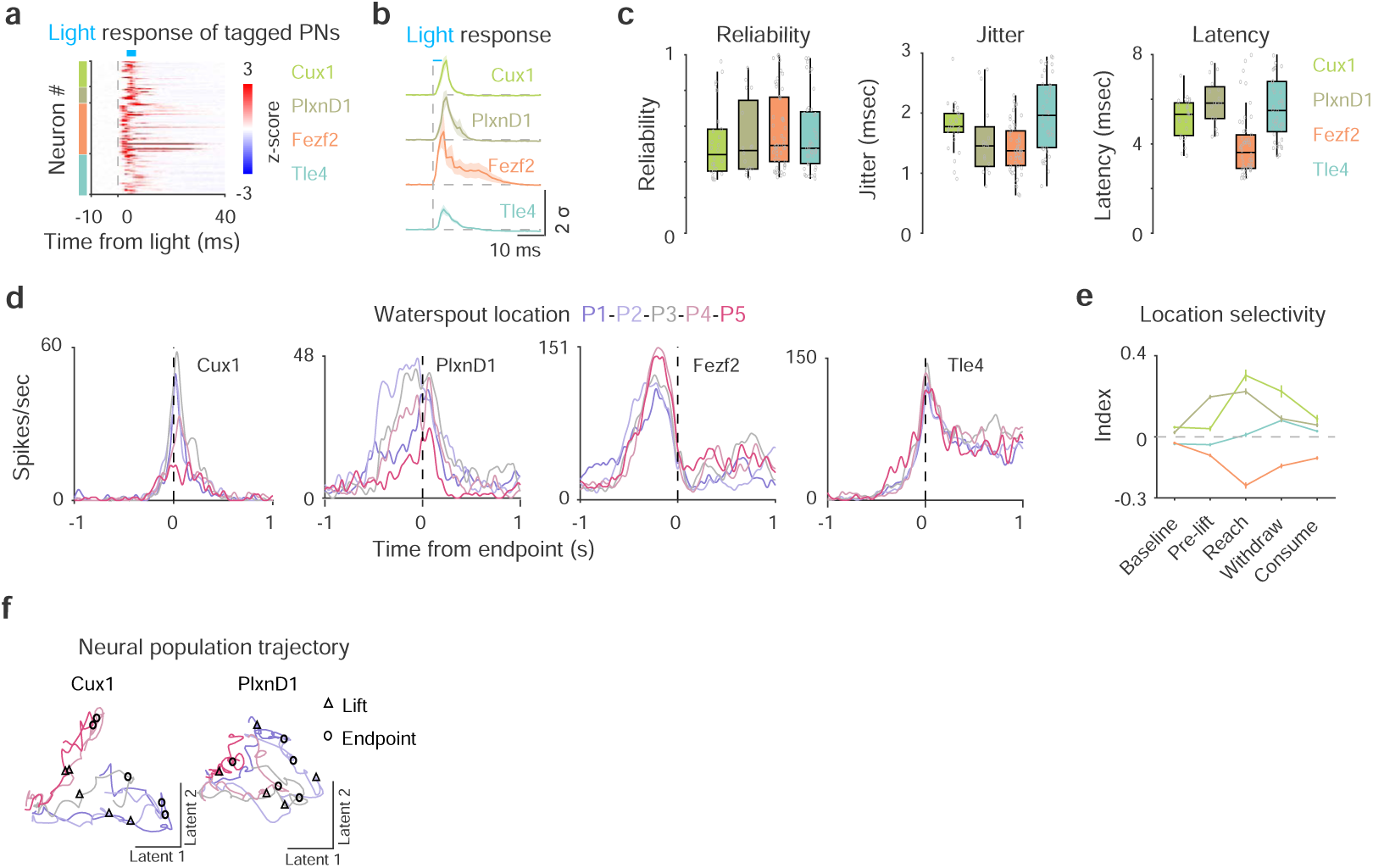
Optogenetic tagging and PN-type specific behavioral correlates. Related to Fig 4. **a.** Light-evoked spiking activity of all tagged PNs. The vertical dashed line indicates light pulse (blue bar) onset. (*n* = 26 IT^Cux1^, 16 IT^PlxnD1^, 50 PT^Fezf2^, and 40 CT^Tle4^ neurons) **b.** Average light-evoked activity of all tagged neurons across different neuron types. The delayed modulation after immediate activity upon light pulses suggests a recurrent connection in PT^Fezf2^. Horizontal dashed lines are 0 reference. **c.** Boxplots of reliability, jitter, and latency of light-evoked spikes of different PN types. **d.** Activity tuning by target location from example IT^Cux1^, IT^PlxnD1^, PT^Fezf2^, and CT^Tle4^ neurons. Vertical dashed lines: advance endpoint. **e.** Waterspout target modulation index of distinct PN types across different action phases. **f.** Average population neural trajectories along the first two principal dimensions for IT^Cux1^ and IT^PlxnD1^.

**Fig S7.**
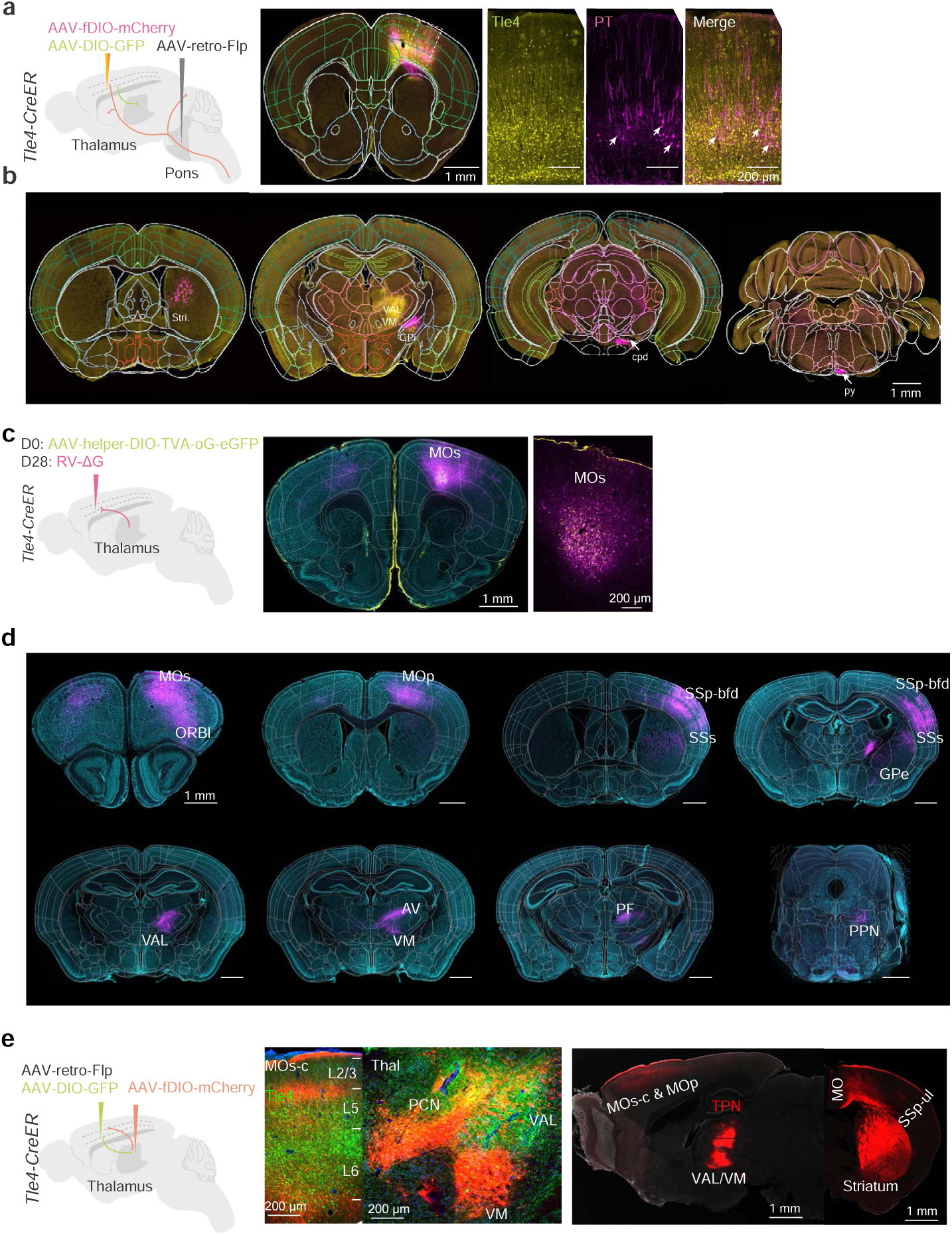
Anatomical connections of MOs-c CT^Tle4^. Related to Fig 5. **a.** Comparison of CT^Tle4^ neurons (yellow) and pons-projecting PT neurons (magenta) in MOs-c. Only 8/144 red neurons show green fluorescence. Arrows indicate PT neurons. **b.** Coronal brain sections show MOs-c CT^Tle4^ neurons (green) project axons exclusively to thalamus without terminals in midbrain or pons, while pons-projecting PT neurons (red) branches in brainstem with few terminals in thalamus. **c.** Left, schematic for mono-synaptic input mapping of MOs-c CT^Tle4^ neurons with rabies tracing (RV). D, Day. The right two panels indicate the EGFP-expressing (yellow) and RV-expressing (magenta) neurons in the MOs-c injection site. Tle4 starters are labeled by both colors (white). Cyan, Nissl stain. **d.** Coronal brain sections show presynaptic partners of MOs-c CT^Tle4^ in orbital frontal cortex (ORBl), sensorimotor cortex (MOp, SSp, SSs), thalamus (VAL, VM, PF, AV) and midbrain (PPN). **e.** Combined retrograde (from MOs-c) and anterograde (from thalamus) tracing (schematic) revealed the recurrent connections between MOs-c and VAL/VM. A mix of AAV with Flp recombinase (*AAV-retro-Flp*) and Cre recombinase dependent GFP (*AAV-DIO-EGFP*) is injected into the MOs-c of *Tle4-CreER* mice. Flp dependent mCherry (*AAV-fDIO-mCherry*) is injected into the thalamus to express in TPN^MOs-c^. MOs-projecting TPNs also send branches to MOp and SSp-ul.

**Fig S8.**
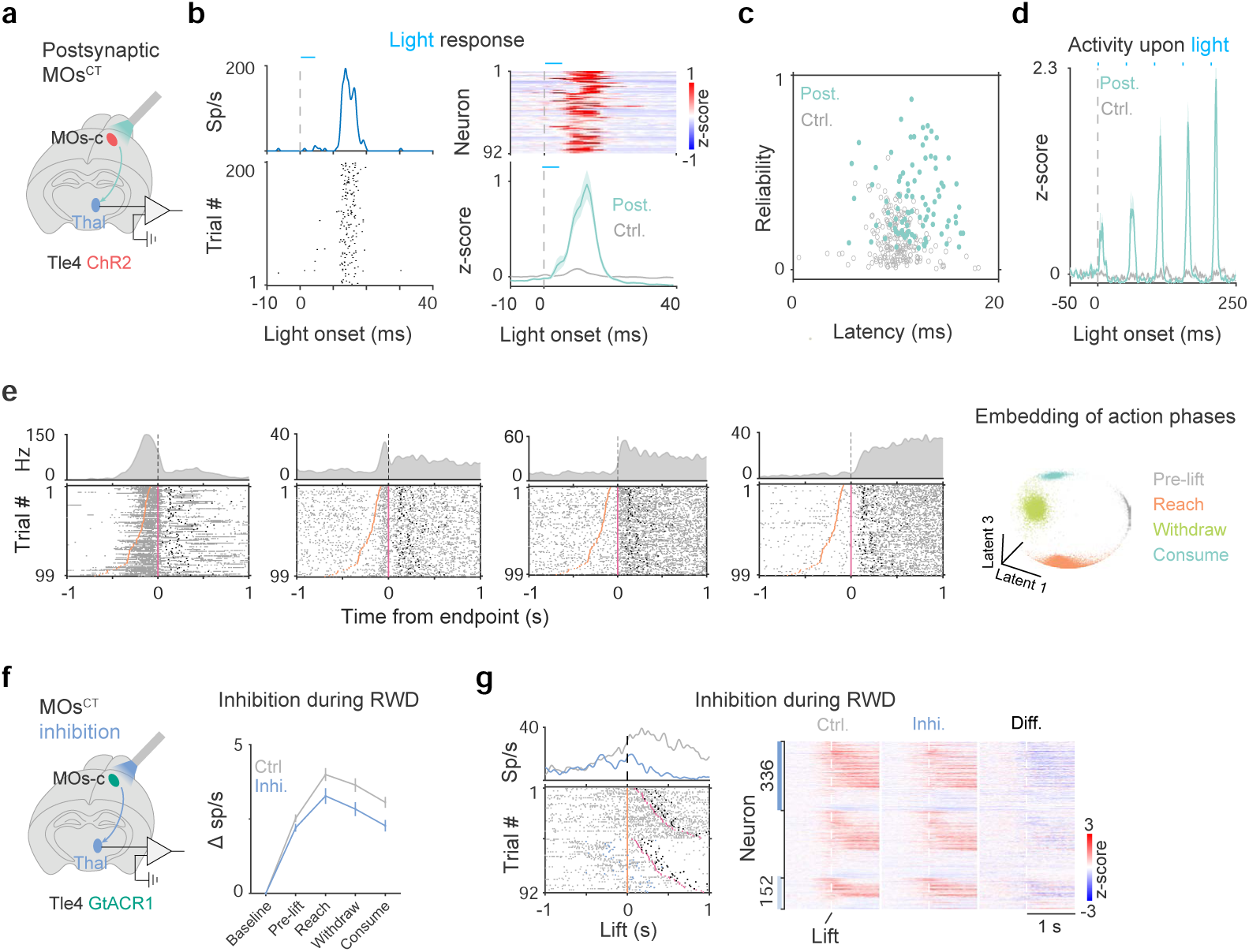
MOs-c CT^Tle4^ modulates thalamic dynamics. Related to Fig 6. **a.** Optogenetic tagging of postsynaptic neurons of MOs-c CT^Tle4^ in thalamus. **b.** Left, light-evoked PETH (top) and raster (bottom) activity of a MOs-c CT^Tle4^ postsynaptic neuron in thalamus (TPN^Tle4-post^). Right, light response of 92 identified TPN^Tle4-post^ (blue) compared with 210 not significantly modulated (gray). **c.** Light-evoked spiking reliability and latency of all thalamic neurons. The same set of neurons were used for the following panels. (*n* = 92 TPN^Tle4-post^ and 210 control (gray circles)) **d.** Facilitation of evoked activity of TPN^Tle4-post^ indicated by a three-fold increase of neural activity upon the 5^th^ light pulse (2.17) compared with that of the 1^st^ one (0.57) with 20 Hz stimulation. **e.** Action phase encoding in simultaneously recorded individual and population neurons in VAL/VM. Raster panels show spikes of individual neurons in relation to actions. Right most panel shows action phase embedding in the latent space of population neural activity revealed by supervised CEBRA model on an example session (*n* = 213 neurons). Each dot represents a time point (1/240 s). Superimposed colors indicate different action phases. **f.** Left, recording thalamic neurons with MOs-c CT^Tle4^ inhibition. Right, average inhibition effect on all thalamus neurons across RWD phases. (*n* = 814 neurons) **g.** Thalamic neuron activity difference between optogenetic inhibition and control conditions. Left, spike raster of a Group I neuron during RWD. Inhibition light was delivered around the lift in random half of the trials. Trials are grouped by inhibition light (bottom) and sorted by the duration between hand lift (orange ticks) and the advance endpoint (pink ticks). Black ticks, first hand licks to consume water. Right, heatmap showing the difference. (*n* = 814 neurons)

**Fig S9.**
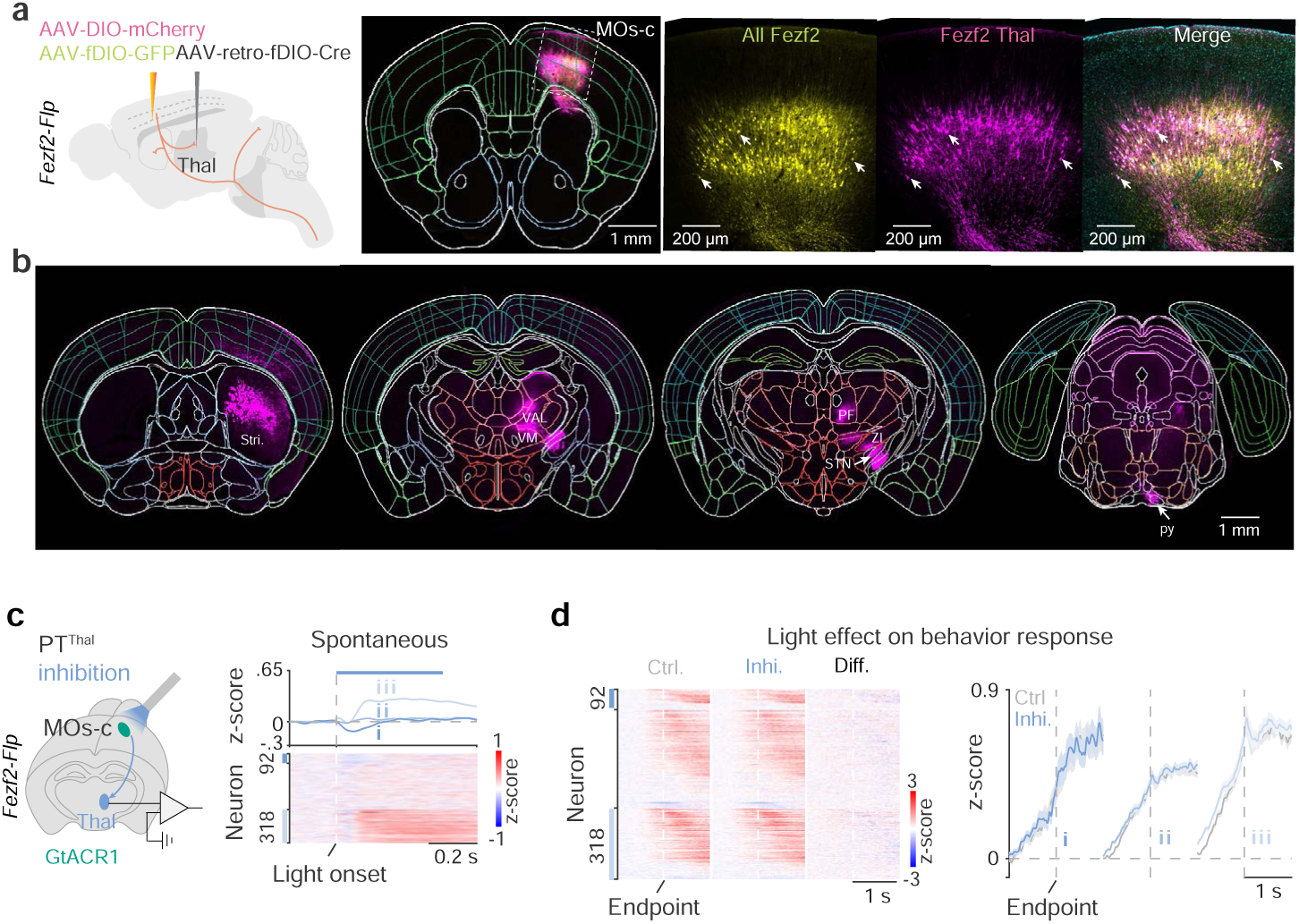
Effects of MOs-c PT^Thal^ on thalamic dynamics. Related to Fig 6. **a.** Two-virus strategy (schematic) to label thalamus-branching PT neurons (PT^Thal^, magenta). *AAV-retro-fDIO-Cre* is injected into the VAL/VM of *Fezf2-Flp* mice. A mix of Cre dependent mCherry (AAV-DIO-mCherry) and Flp recombinase dependent GFP (*AAV-fDIO-EGFP*) is injected to the MOs-c. This strategy indicates ∼40% (545/1334 neurons) of Fezf2 neurons branch in thalamus (*n* = 2 mice). Arrows indicate double-labeled neurons. Scale, 1 mm for the whole section and 200 μm for zoom-in view. **b.** Coronal brain sections show MOs-c PT^Thal^ neurons (magenta) collaterals in the striatum (Stri.), STN, ZI, and other midbrain and brainstem nuclei besides VAL/VM. Scale, 1 mm. **c.** Electrophysiological recording of thalamus neurons upon optogenetic inhibition of thalamus-branching PT (PT^Thal^) in MOs-c. Left, expressing GtACR1 in MOs-c PT^Thal^ neurons with the two-virus strategy. Right, average effect of MOs-c PT^Thal^ on the spontaneous firing of 814 thalamic neurons. Neurons were divided into 92 decreased Group I, 439 non-modulated Group II, and 318 increased Group III. **d.** Thalamic activity difference between MOs-c PT^Thal^ inhibition and control conditions. (*n* = 849 neurons)

## Notes

### Competing Interest Statement

The authors have declared no competing interest.

### Summary of Updates

Figure pannels rearranged; New data added.

